# A Programmable Ontology Encompassing the Functional Logic of the *Drosophila* Brain

**DOI:** 10.1101/2021.12.28.474399

**Authors:** Aurel A. Lazar, Mehmet Kerem Turkcan, Yiyin Zhou

## Abstract

The *Drosophila* brain has only a fraction of the number of neurons of higher organisms such as mice and humans. Yet the sheer complexity of its neural circuits recently revealed by large connectomics datasets suggests that computationally modeling the function of fruit fly brain circuits at this scale posits significant challenges.

To address these challenges, we present here a programmable ontology that expands the scope of the current *Drosophila* brain anatomy ontologies to encompass the functional logic of the fly brain. The programmable ontology provides a language not only for modeling circuit motifs but also for programmatically exploring their functional logic. To achieve this goal, we tightly integrated the programmable ontology with the workflow of the interactive FlyBrainLab computing platform. As part of the programmable ontology, we developed NeuroNLP++, a web application that supports free-form English queries for constructing functional brain circuits fully anchored on the available connectome/synaptome datasets, and the published worldwide literature.

In addition, we present a methodology for including a model of the space of odorants into the programmable ontology, and for modeling olfactory sensory circuits of the antenna of the fruit fly brain that detect odorant sources. Furthermore, we describe a methodology for modeling the functional logic of the antennal lobe circuit consisting of massive local feedback loops, a characteristic feature observed across *Drosophila* brain regions. Finally, using a circuit library, we demonstrate the power of our methodology for interactively exploring the functional logic of the massive number of feedback loops in the antennal lobe.

## 1 Introduction

### 1.1 Challenges in Discovering the Functional Logic of Brain Circuits in the Connectomic/Synaptomic Era

Large scale foundational surveys of the anatomical, physiological and genomic architecture of brains of mice, primates and humans have shown the enormous variety of cell-types [1, 2, 3], diverse connectivity patterns with fan-ins and fan-outs in the tens of thousands and extensive feedback that vary both within and between brain regions [4]. The last decade also saw an exponential growth in neuroscience data gathering, collection and availability, starting with the cubic millimeter brain tissue in mice and humans [5]. However, due to the sheer magnitude and complexity of brains of higher organisms, even with such data at hand, we are far behind in our understanding of the principles of neural computation in the brain.

Prior studies have highlighted the need for developing means of formally specifying and generating executable models of circuits that incorporate various types of brain data, including the heterogeneity and connectivity of different cells types and brain circuits, neurophysiology recordings as well as gene expression data. In principle, a whole brain simulation can be instantiated by modeling all the neurons and synapses of the connectome/synaptome with simple dynamics such as integrate-and-fire neurons and *α*-synapses, with parameters tuned according to certain criteria [6]. Such an effort, however, may fall short of revealing the fundamental computational units required for understanding the functional logic of the brain, as the details of the units of computation are likely buried in the uniform treatment of the vast number of neurons and their connection patterns.

It is, therefore, imperative to develop a formal reasoning framework of the functional logic of brain circuits that goes beyond simple instantiations of flows on graphs generated from the connectome. A framework is needed for building a functional brain from components whose functional logic can be readily envisioned, and for exploring the computational principles underlying these components given the available data.

Recently released connectome, synaptome and transcriptome datasets of the *Drosophila* brain and ventral nerve cord (VNC) presents a refreshing view for the study of neural computation [7, 8, 9]. These datasets present challenges and opportunities for hypothesizing and uncovering the fundamental computational units and their interactions.

### 1.2 Modeling the Functional Logic of Fruit Fly Brain Circuits with Feedback Loops

The fruit fly brain can be subdivided into some 40 neuropils. The concept of local processing units (LPU) was introduced in the early works of the fly connectome to represent functional subdivisions of the fruit fly brain circuit architecture [10]. LPUs are characterized by unique populations of local neurons whose processes are restricted to specific neuropils.

It was not until the release of follow up electron microscopy (EM) connectome datasets that the minute details of the connectivity of these local neurons were revealed [8, 7, 11, 12]. Often times, local neurons within each neuropil form intricate feedback circuits with a massive number of feedback loops.

For example, the antennal lobe of the early olfactory system, consists of the axons of olfactory sensory neurons (OSNs) as inputs (depicted in Figure 1A in darker colors), the antennal lobe projection neurons (PNs) as outputs (in Figure 1A in brighter colors), and a large collection of local neurons (in Figure 1A in transparent white). The adjacency matrix of the connectivity graph of the AL circuit is shown in Figure 1B.

**Figure 1:**
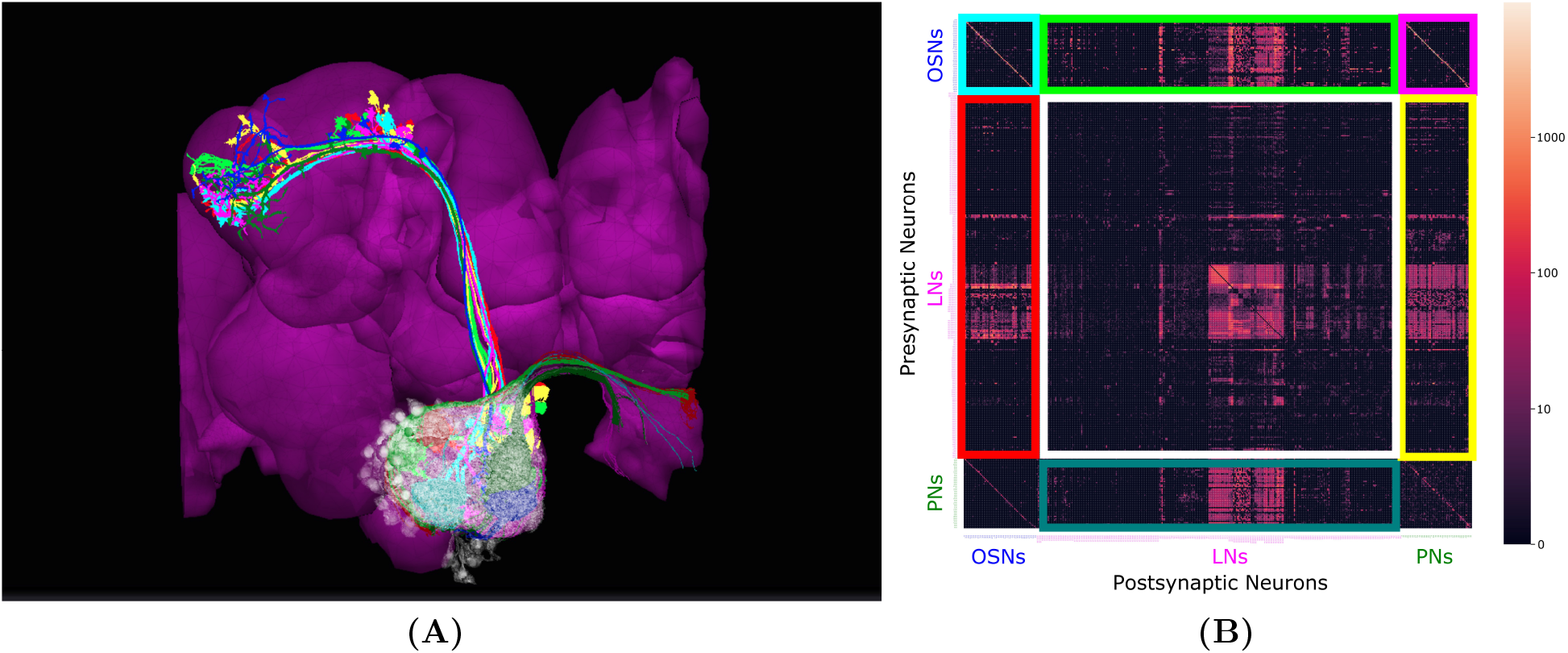
Massive Number of Feedback Loops in the Antennal Lobe. (A) Antennal Lobe circuit involving OSNs (darker colors), PNs (brighter colors) and LNs (transparent white). Select OSNs, PNs and LNs are shown. (B) The adjacency matrix of the connectivity graph of the neurons in the AL, with all OSNs expressing the same OR merged into a single neuron group node, and all PNs in the same glomerulus merged into a single neuron group node. Matrix elements indicate the number of synapses from a presynaptic neuron (or neuron group) to a postsynaptic neuron (or neuron group). Magenta block on top right: submatrix of the feedforward connectivity from OSNs to PNs in each glomerulus. Green block on the top: submatrix of the feedforward connectivity from OSNs to LNs. Blue block on the bottom: submatrix of the connectivity from PNs to LNs. Red block on the left: submatrix of the feedback connectivity from LNs to OSNs. Yellow block on the right: submatrix of the feedback connectivity from LNs to PNs. White block in the middle: submatrix of the connectivity between LNs.

The axons of the OSNs expressing the same olfactory receptor (OR) project into the same glomerulus where they provide inputs to uniglomerular PNs (uPNs) whose dendrites only extend within the same glomerulus. Such connections form the feedforward signaling path in the antennal lobe (see the magenta-colored block in Figure 1(B).

While not all neuropils share such glomerular structure, two more features in the AL connectivity patterns can be found in many other neuropils.

First, OSNs expressing the same OR exhibit strong axon-axonal connections but not with OSNs expressing other ORs (see the cyan-colored block corresponding to the OSN-to-OSN connectivity on the top left of Figure 1(B)). Similar axonal connections can be observed between Kenyon Cells (KCs) of the mushroom body (MB) [7], between Lobular Columnar (LC) neurons in the optic glomeruli (OG) [8], and between the ring neurons of the ellipsoid body (EB) [13].

Second, local neurons receive inputs from OSNs and PNs (see green and blue blocks, corresponding to OSN-to-LN and PN-to-LN connectivity, respectively, in Figure 1(B)). They also provide feedback to OSNs and PNs (see red and yellow blocks, respectively, in Figure 1(B)). In addition, LNs also synapse onto other LNs (white block in Figure 1(B)). Given the simplicity of the feedforward signaling path and the complex nature of feedback driven by LN connectivity, these massive number of feedback pathways must underlie the functional logic of the AL circuit.

Massive feedback loops can be ubiquitously found across other brain regions, for example in the medulla [12], lateral horn, mushroom body [8], central complex [13], etc. Since the AL has a connectivity structure that in many ways is representative, for simplicity and clarity in the rest of this work we will be mostly focused on characterizing the AL circuit.

Finally, note that in mammals, particularly in the visual system, feedback pathways have long been considered to be a key component of the architecture of brain circuits [14]. Yet, the lack of detailed brain circuit connectivity in these higher organisms has not yet provided much insight into the functional role played by the feedback circuits. The connectome/synaptome of the fruit fly opens new avenues for discovering the full complexity and computational principles underlying feedback circuits.

### 1.3 A Programmable Ontology Encompassing the Functional Logic of the Fruit Fly Brain Circuits

Traditionally, ontologies formally define the classification of the anatomical structure of the *Drosophila* nervous system and the ownership relationships among anatomical entities [15, 16]. However, existing ontologies lack computational primitives/motifs, such as feedback pathways that can be more readily associated with the functional role of brain circuits. Furthermore, characterizing the functional logic of sensory circuits calls for modeling the environment the fruit flies live in. The space of natural sensory stimuli that the fruit flies constantly sample has not been discussed in the formal ontology of the fly brain anatomy. It is often neglected in the neuroscience literature, but essential in defining, characterizing and evaluating the functional logic of its brain circuits.

As we argued here, expanding the scope of the classical ontology to encompass the natural sensory stimuli and the functional logic of the *Drosophila* brain bridges the gap between the two fields and greatly benefits both. Such a *programmable ontology* will provide a language not only for describing but also for executing the functional modules of, for example, the massive number of feedback loops observed in brain circuits and help make their contribution to brain function transparent.

The proposed programmable ontology is tightly integrated with the workflow of the interactive FlyBrainLab [16] computing platform, as elaborated in Figure 2. The workflow in Figure 2 consists of 3 steps. First, 3D visualization (see Materials and Methods) of fly brain morphology data is explored and candidate anatomical structures defining functional units and modules (Figure 2 left) identified. Second, the candidate biological circuits are mapped into executable circuits that provide an abstract representation of the circuit in machine language (Figure 2 middle). Third, the devised executable circuits are instantiated for the interactive exploration of their functional logic with a highly intuitive graphical interface for configuring, composing and executing neural circuit models (Figure 2 right, see Materials and Methods).

**Figure 2:**
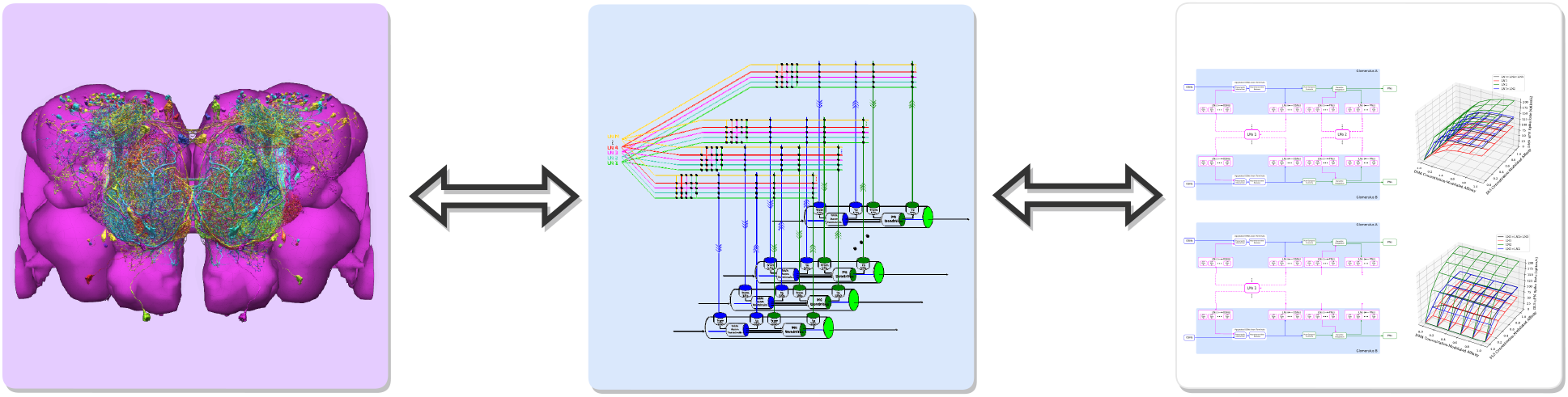
The workflow of discovery of the functional logic of the fruit fly brain. Left: 3D visualization and exploration of fly brain data. Middle: Creation of executable circuits. Right: Interactive exploration of the functional logic of executable circuits.

The main rationale for the tight integration of the programmable ontology into the FlyBrainLab workflow of discovery is to fully anchor it onto biological data and the worldwide literature that describes it. FlyBrainLab fully supports the programmability of the ontology while easily supporting various computational schemes used for interrogating the functional logic of brain circuits.

## 2 Exploring the Morphology of Cell-Types and Feedback Circuits

Recent releases of large-scale connectomic/synaptomic datasets have enabled experimental and computational neuroscientists to explore neural circuits in unprecedented detail. As Figure 2 suggests, understanding the functional logic of fruit fly brain circuits starts with the exploration of fly brain connectome/synaptome datasets. To efficiently explore these datasets requires, however, knowledge of both the biological nomenclature and programming tools. These skills are often limited to members of their respective communities, such as neurobiologists and computer scientists.

To close the programming gap, we developed the natural language query interface NeuroNLP to support highly sophisticated English queries of *Drosophila* brain datasets, including morphology and position of neurons (cell-type map), connectivity between neurons (connectome) and distribution and type of synapses (synaptome) [17, 16]. Moreover, NeuroNLP provided the first open neurophysiology data service for the fruit fly brain (activity map). However, the NeuroNLP rule-based query engine could only map pre-designed sentence structures into NeuroArch Database [18] queries, thereby limiting its usage. In particular, users unfamiliar with the nomenclature used in a dataset may have found it difficult to query for particular neuron types.

In this section, we introduce NeuroNLP++, a substantially upgraded NeuroNLP web application, that alleviates these limitations and helps users to explore fruit fly brain datasets with free-form English queries. In Section 2.1, we introduce the NeuroBOT natural language engine supporting NeuroNLP++ and describe its usage in querying cell-types. In Section 2.2, we demonstrate how to use NeuroNLP++ to query feedback circuits.

### 2.1 Exploring the Morphology of Cell-Types with NeuroNLP++

Expanding upon the NeuroNLP query interface of the Fruit Fly Brain Observatory (FFBO), NeuroNLP++ provides two additional key advances. First, NeuroNLP++ interprets and answers free-form English queries that are well beyond the natural language capabilities of NeuroNLP. Second, NeuroNLP++ not only visualizes neuron/synapses but also links them to the worldwide fruit fly brain literature.

To interpret free-form English queries, we built the DrosoBOT engine that can be accessed through the NeuroNLP++ application (see Materials and Methods).

DrosoBOT associates descriptive terms of neurons in the fruit fly brain with connectomic datasets. For example, it integrates cell-types or lineages from the Drosophila Anatomy Ontology (DAO) [15] and matches them against neurons in the Hemibrain connectome dataset [8]. Given a query, DrosoBOT employs state-of-the-art document retrieval techniques [19] to find relevant descriptions. Simple examples include “what types of local neurons are in the antennal lobe?” or “which neurons are known to respond to carbon dioxide?”.

In what follows we give some examples of DrosoBOT query results. A result of a DrosoBOT query is a list of the most relevant cell-types displayed in the Info Panel of the NeuroNLP++ user interface, as shown in Figure 3(B). Each entry lists the name of the cell-type, a link to the DAO, as well as a description of the cell-type. It also includes a UI button for adding to the workspace the neurons associated with the entry.

**Figure 3:**
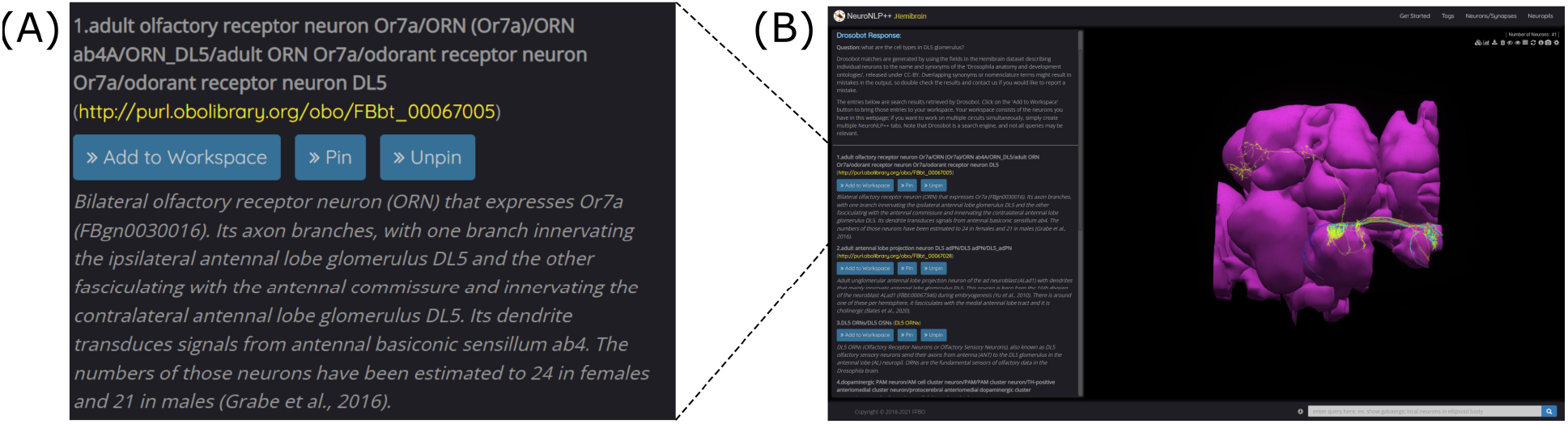
NeuroNLP++ user interface and query results from DrosoBOT. (A) A typical entry of Drosobot query results displayed in the Info Panel of the NeuroNLP++ application in response to a ”what are the cell-types in the DL5 glomerulus?” query. (B) The user interface of NeuroNLP++ with the Info Panel on the left and 3D visualization workspace on the right.

In the first example, we started by asking “what are the cell-types in the DL5 glomerulus”. Figure 3(A) shows one of the query results, including names and synonyms of the OSNs that project into the DL5 glomerulus, as well as the ontological description of these OSNs along with specific entries of the relevant literature. In addition to the OSNs, a second cell type is the DL5 adPN. We add both of these cell-types into the visualization workspace with different colors (see Figure 4(A), right). For improved visualization, NeuroNLP++ provides a graph view of current neurons in the workspace at single-cell or cell-type level (see Materials and Methods). The cell-type graph view of the neurons in the DL5 glomerulus is depicted on the left in Figure 4(A). Here, the red and the yellow nodes represent the OSNs and PNs, respectively, and the arrow from the red to the yellow node indicates that the OSNs provide inputs to the PNs. The colors in the graph view match those in the 3D morphology visualization. The graph view is also interactive, allowing users to highlight the corresponding neurons in the 3D visualization.

**Figure 4:**
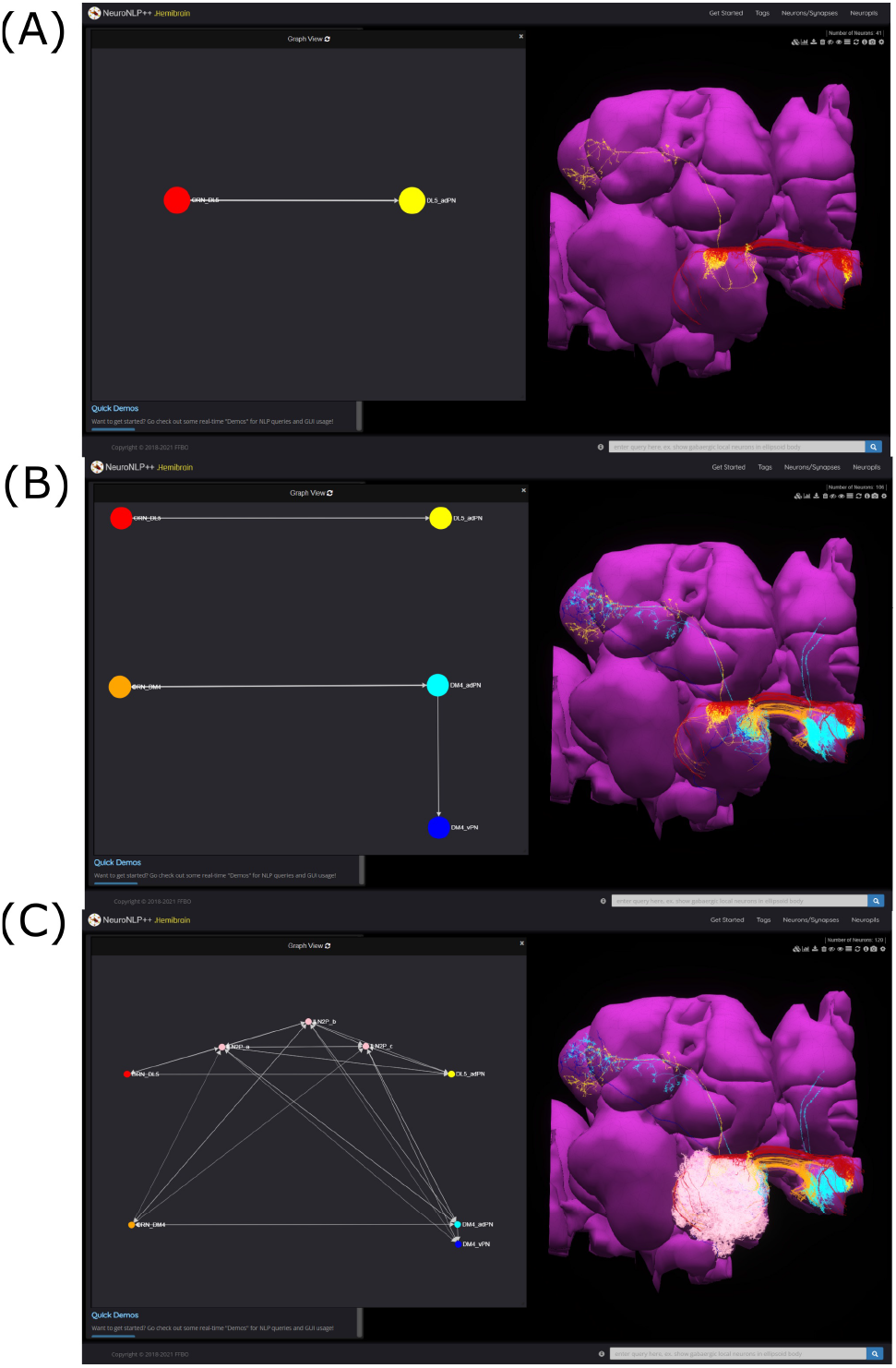
Free-form English queries of the AL with NeuroNLP++. (A) The result to the query “what are the cell types in the DL5 glomerulus?”, consisting of the OSNs that project into the DL5 glomerulus (red) and the PNs with dendrites in the DL5 glomerulus (yellow). (left) Cell-type connectivity graph of the visualized neurons. (right) Morphology of the retrieved neurons. (B) The result to the query “what are the cell types in the DM4 glomerulus?”, in addition to (A), consisting of the OSNs (orange) projecting into the DM4 glomerulus and the adPNs (cyan) and vPNs (blue) with dendrites in the DM4 glomerulus. (left) Cell-type level connectivity graph of the resulting neurons. (right) Morphology of the retrieved neurons. (C) Results to the query “what are the patchy local neurons?”, in addition to (B). The resulting LNs are in pink. (left) Cell-type connectivity graph of the visualized neurons. (right) Morphology of the retrieved neurons.

We then asked “what are the cell-types in the DM4 glomerulus”. The resulting neurons are added to the workspace and their cell-type graph is depicted in Figure 4(B). These include the OSNs (orange) that project into the DM4 glomerulus and two types of PNs, namely adPNs (cyan) that project to both the MB and LH, and vPNs (blue) that only project to the LH. The graph also confirms that the two glomeruli run in parallel. Finally, we asked “what are the patchy local neurons?”. The resulting neurons are shown in white in Figure 4(C), together with the cell-type graph of the entire circuit. The connectivity graph suggests the presence of strong feedback components within and between the two otherwise disjoint glomerular circuits.

To explore the diversity of LNs in the AL, we needed to classify cell-types based on their morphology. We launched the query “what are the types of local neurons in the antennal lobe?”. DrosoBOT provides a complete list of the currently known LN types in the AL. In Figure 15 in the Appendix A, we list all these LN types, the number of neurons of each type, and an example morphology of neuron type. The morphology of the neurons is colored by their glomerular arborization.

Finally, we mention here queries of neurons that may be located downstream of the AL. We asked “what dopaminergic neuron subtypes are there” (Figure 5(A)), and “what cell types are there in lateral horn” (Figure 5(B)). The query results revealed a variety of cell types. These may provide a starting point for exploring novel cell-types associated with other neuropils and can guide additional rule-based queries.

**Figure 5:**
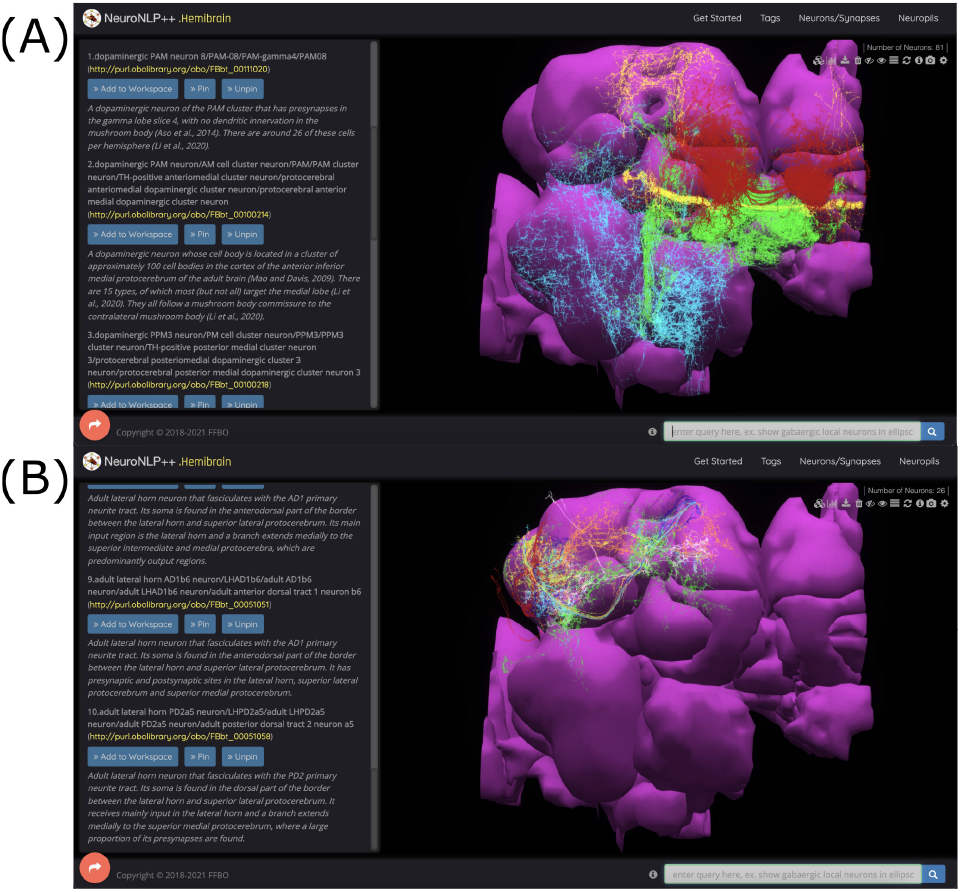
Querying cell-types downstream of the AL with NeuroNLP++. (A) The result to the query “what dopaminergic neuron subtypes are there?” with the first few entries added to the workspace. (red) PAM08 neurons, (green) PPM3 neurons, (yellow) PAM09 neurons, (cyan) PPM01 neurons. (B) The result to the query “what cell types are there in lateral horn?” with the first few entries added to the workspace. (red) AV4b3 neurons, (green) LH centrifugal neurons, (yellow) AD1c3 neurons, (cyan) AD1c2 neurons, (blue) AD1d2 neurons, (orange) AD1b6, (white) PD2a5 neurons.

### 2.2 Exploring Massive Feedback Circuits with NeuroNLP++

As discussed above, massive feedback circuits are major targets of the study of the functional logic in the fruit fly brain. Therefore, in addition to querying cell-types, we also built into NeuroNLP++ capabilities to query predefined circuit motifs including those that exhibit feedback loops.

Different feedback circuits can be constructed as ontological entities loaded into DrosoBOT, and queried with NeuroNLP++. We precomputed different types of feedback circuits for each glomerulus (see Materials and Methods) and considered a number of specific feedback loops consisting of local neurons. For example, the circuit consisting of LNs that receive inputs from and provide feedback to OSNs but has no interaction with PNs is named an LN1-type feedback loop. Similarly, the circuit consisting of LNs that receive inputs from and provide feedback to PNs but has no interaction with OSNs is named an LN2-type feedback loop.

To query for feedback loops associated with OSNs and PNs in the DL5 glomerulus, *i.e.*, starting from Figure 4(A), we requested: “show available feedback loops”. Figure 6 depicts two different types of feedback loops in response to this query. Notably, the Info Panel explicitly lists the different types of feedback loops. In Figure 6(A), we added the 19 LNs (pink) that form the LN2 feedback loop. From the graph view, we confirm that these feedback loops are only associated with the DL5 PNs (yellow) but not OSNs (red) node. The arrows into the adPN node indicate the feedback pathway from LNs into the adPN. In Figure 6(B), we added the 26 LNs that form feedback loops that exhibit both LN1- and LN2-type characteristics. We can observe from the graph view that these LNs (pink) form feedback loops with both OSNs (red) and PNs (yellow) nodes.

**Figure 6:**
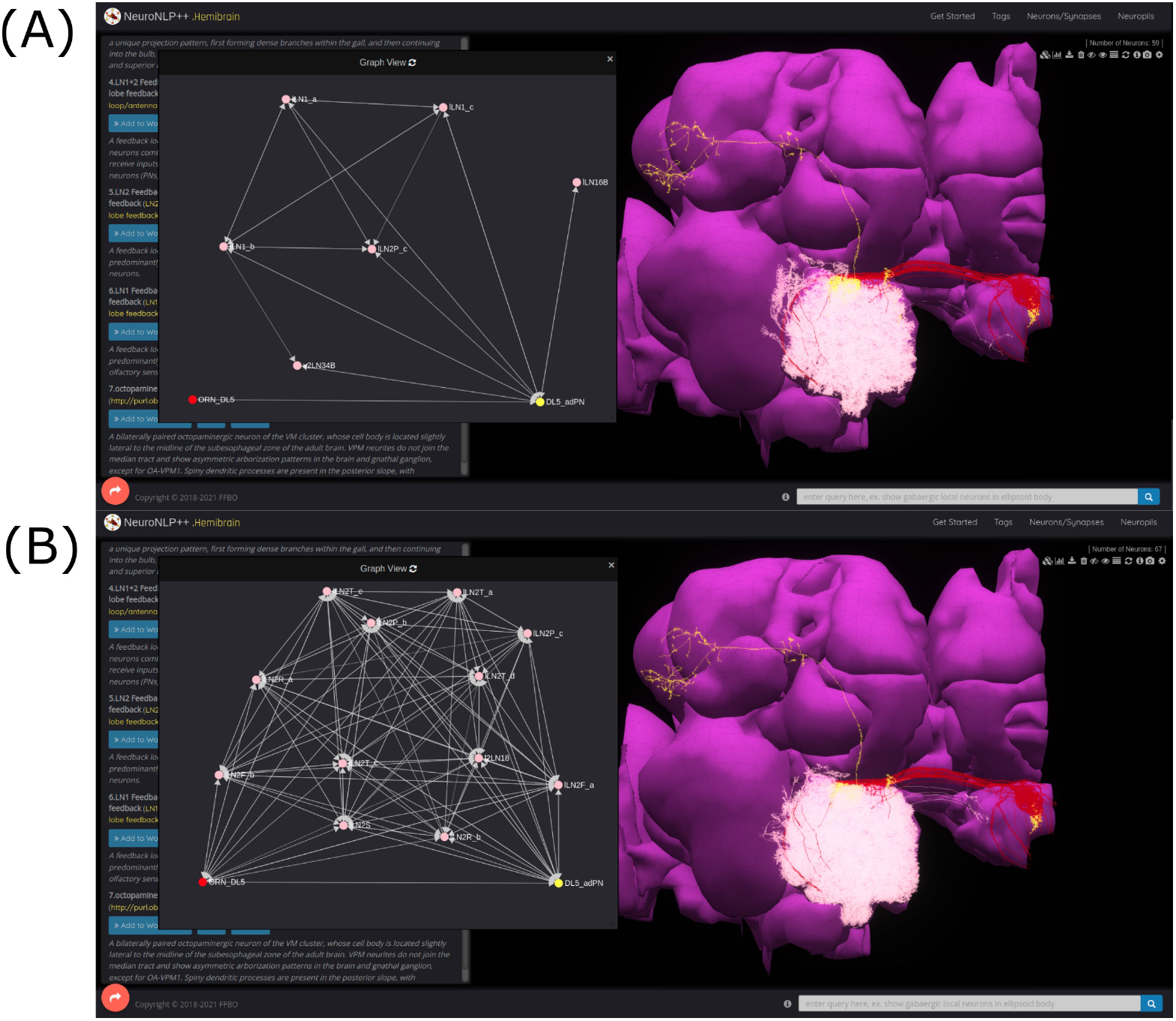
Exploring feedback circuits with NeuroNLP++. The result to the query “show available feedback loop” when starting from Figure 4(A) with OSNs that project into the DL5 glomerulus and PNs with dendrites in DL5 glomerulus. (A) LN2-type feedback loop (receives inputs from and only feeds back to PNs). (red) OSNs. (yellow) PNs. (pink) LNs. (B) Feedback loops that exhibit both LN1- and LN2-type characteristics (receive inputs from both OSNs and PNs and provide feedback to them). (red) OSNs. (yellow) PNs. (pink) LNs.

Concluding, DrosoBOT is built to be modular and enables different types of feedback loops to be added to our programmable ontology to facilitate the construction and visualization of circuit motifs [20]. NeuroNLP++ represents a step towards a more intuitive and natural way of extracting information from large connectome/synaptome datasets that are relevant for the in-depth study of the functional logic of brain circuits. In addition, the capability to anchor the queried connectome/synaptome data onto the published worldwide literature provides much needed awareness of the prior existing knowledge regarding these circuits.

## 3 Creating Executable Fruit Fly Brain Circuit Models

In the previous section, by establishing the NeuroNLP++ natural language query interface for exploring the morphology of fruit fly brain circuits, we effectively created an ontology of the fruit fly brain consisting of the existing anatomical ontology, the connectome/synaptome datasets and the published worldwide literature. Moreover, we provided visualization tools for extracting, what are thought to be, functionally significant circuits. Our goal in this section is to demonstrate how this programmable ontology can be extended to encompass the key elements needed for exploring the functional logic of the fruit fly brain circuits.

The significance of modeling the space of stimuli for characterizing the I/O of functional circuits arises throughout the early sensory systems, e.g., in early olfaction, vision, audition, mechanosensation, etc.. The odorant space and the visual field are examples that come to mind. See, for example, [21] and [22].

Given the current datasets, we shall describe here how some of the better characterized neuropils can be modeled in detail. Due to space limitations, we will only present in what follows our methodology of a receptor-centric modeling of the space of odorant stimuli, as well as modeling of the olfactory processing in the antenna and the antennal lobe of the fruit fly brain. How to apply the same methodology to other neuropils of the fruit fly brain will be presented elsewhere.

### 3.1 Receptor-Centric Modeling the Space of Odorant Stimuli

To fully characterize the functional logic of a sensory circuit calls for modeling the environment the studied organism lives in, a rather difficult undertaking. To model the environment, we first have to define the *space of stimuli*. The space of stimuli has never been discussed in the context of a formal ontology of the fruit fly brain anatomy. It is often neglected in the neuroscience literature, but essential in defining, characterizing and evaluating the functional logic of brain circuits.

The Chemical Abstracts Service (CAS) registry has currently 156 million organic and inorganic substances registered [23]. Distinguishing between odorants in the CAS registry seems to be a problem of enormous complexity. How does the fly approach this problem? As a first step in the encoding process, the odorant receptors bind to the odorants present in the environment and that are of interest to the fly. The fruit fly has some 51 receptors whose binding and dissociation from odorant molecules characterizes their identity. In addition to odorant identity, the odorant concentration amplitude is another key feature of the odorant space.

The odorant space considered here consists of pure and odorant mixtures. Pure odorants are mostly used in laboratory settings for studying the capabilities and the function of the early olfactory circuits. Odorant mixtures widely arise in the living environment. Following [21], the identity of an odorant can be modeled by a 3D tensor trio (**b, d, u**). The 3D tensor **b** with entries [**b**]_*ron*_ is called the odorant-receptor binding rate and models the association rate between an odorant *o* and a receptor of type *r* expressed by neuron *n* (see also Figure 7). The 3D tensor **d** with entries [**d**]_*ron*_ denotes the *odorant-receptor dissociation rate* and models the detachment rate between an odorant *o* and a receptor of type *r* expressed by neuron *n* (see also Figure 7). We denote the concentration of odorants as **u**(*t*), where [**u**]_*o*_(*t*) denotes the concentration of odorant *o, o* = 1, 2*, ..., O*. The odorant concentration can be any arbitrary continuous waveform (see also Figure 7). For a pure odorant 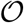, [**u**]_*o*_(*t*) = 0, 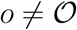. A set of odorant waveforms modeled by the tensor trio (**b, d, u**(*t*)) is graphically depicted in Figure 7. Often, for simplicity, the binding rate [**b**]_*ron*_ and the dissociation rate [**d**]_*ron*_, for a given odorant *o* and a given receptor-type *r*, are assumed to take the same value for all neurons *n* = 1, 2*, ..., N*, expressing the same receptor-type *r*.

**Figure 7:**
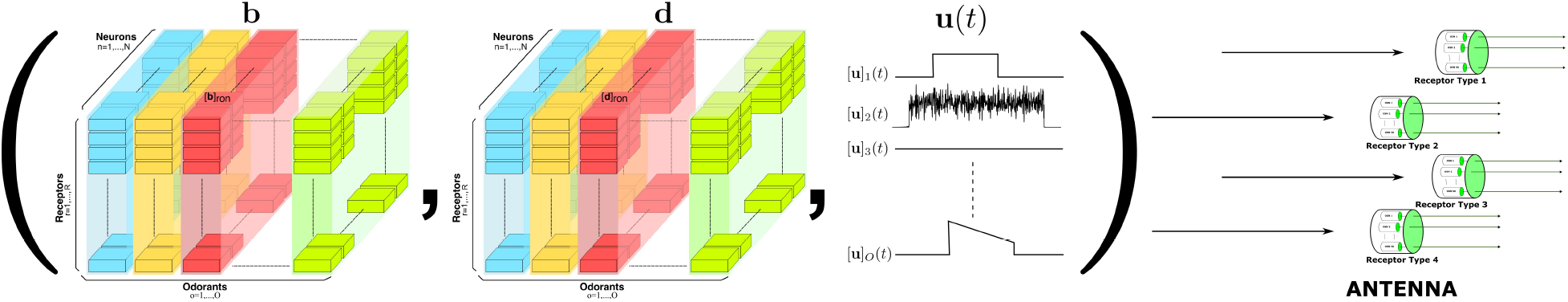
Elements of the odorant space are defined by the odorant-receptor binding rate, dissociation rate and concentration amplitude tensor trio (**b, d, u**(*t*)). For a given neuron *n* = 1, 2*, ..., N*, the binding rate and dissociation rate values are, respectively, denoted by [**b**]_*ron*_ and [**d**]_*ron*_, for all *r* = 1, 2*, ..., R*, and *o* = 1, 2*, ..., O*. The odorants then interact with the receptors expressed by the Olfactory Sensory Neurons in the Antenna (right).

The elements of the odorant space are not defined by the (largely intractable) detailed/- precise chemical structure of the odorants. Rather, they are described by the rate of activation/deactivation between odorants and olfactory receptors. The tensor trio determines what types of sensors (olfactory receptors) will be activated by a certain odorant, and the level of activation will be jointly governed by the identity and the concentration waveform of the odorant. More precisely, the overall activation of the sensors is determined by the value of the odorant-receptor binding rate modulated by the odorant concentration profile [21].

### 3.2 Receptor-Centric Modeling of the Antenna Circuit

The antenna circuit of the early olfactory system of the fruit fly consists of approx. 2,500 parallel Olfactory Sensory Neurons (OSNs) that are randomly distributed across the surface of the maxillary palp and antennae. In what follows, we will refer to the set of all OSNs on one side of the fruit fly brain as an antenna/maxillary palp (ANT) local processing unit (LPU).

The OSNs, depicted in Figure 7(right) in groups based on the olfactory receptors (ORs) that they express, form parallel circuits. For simplicity, we assumed that the number of OSNs expressing the same receptor-type is *N*.

For each OSN, the odorants are first transduced by an olfactory transduction process (OTP) that depends on the receptor-type [21]. Each of the resulting transduction currents drive biophysical spike generators (BSGs) that produce spikes at the outputs of the antennae (see Materials and Methods). Note that unlike the OTP whose I/O characterization depends on the receptor-type, the BSGs of OSNs expressing different receptor-types are assumed to be the same.

### 3.3 Modeling and Constructing the Massive Feedback Circuits of the Antennal Lobe

The overall goal of this section is to develop a methodology for modeling and constructing circuits of arbitrary complexity of the Antennal Lobe. The methodology demonstrated here is generalizable to the other neuropils in the early olfactory system of the fruit fly brain, including the mushroom body and the lateral horn.

#### Modeling Glomerular Feedforward Circuits of the Antennal Lobe

As sketched in Figure 4, the AL exhibits a glomerular structure. Each glomerulus is primarily driven by the feedforward connections between the OSNs expressing the same OR and the corresponding PNs.

To model a glomerulus, we create a circuit diagram as depicted in Figure 8. We abstract the group of OSNs that project into the glomerulus as a single OSN (cell-type). Similarly, the group of PNs projecting into the same glomerulus is abstracted as a single PN (cell-type). We only consider here the PNs that send their axons to both the MB and LH, and, thereby, primarily omit the vPNs that only output to the LH but not the MB. As shown in Figure 4(D), these PNs typically receive inputs from the other PNs rather than OSNs.

**Figure 8:**
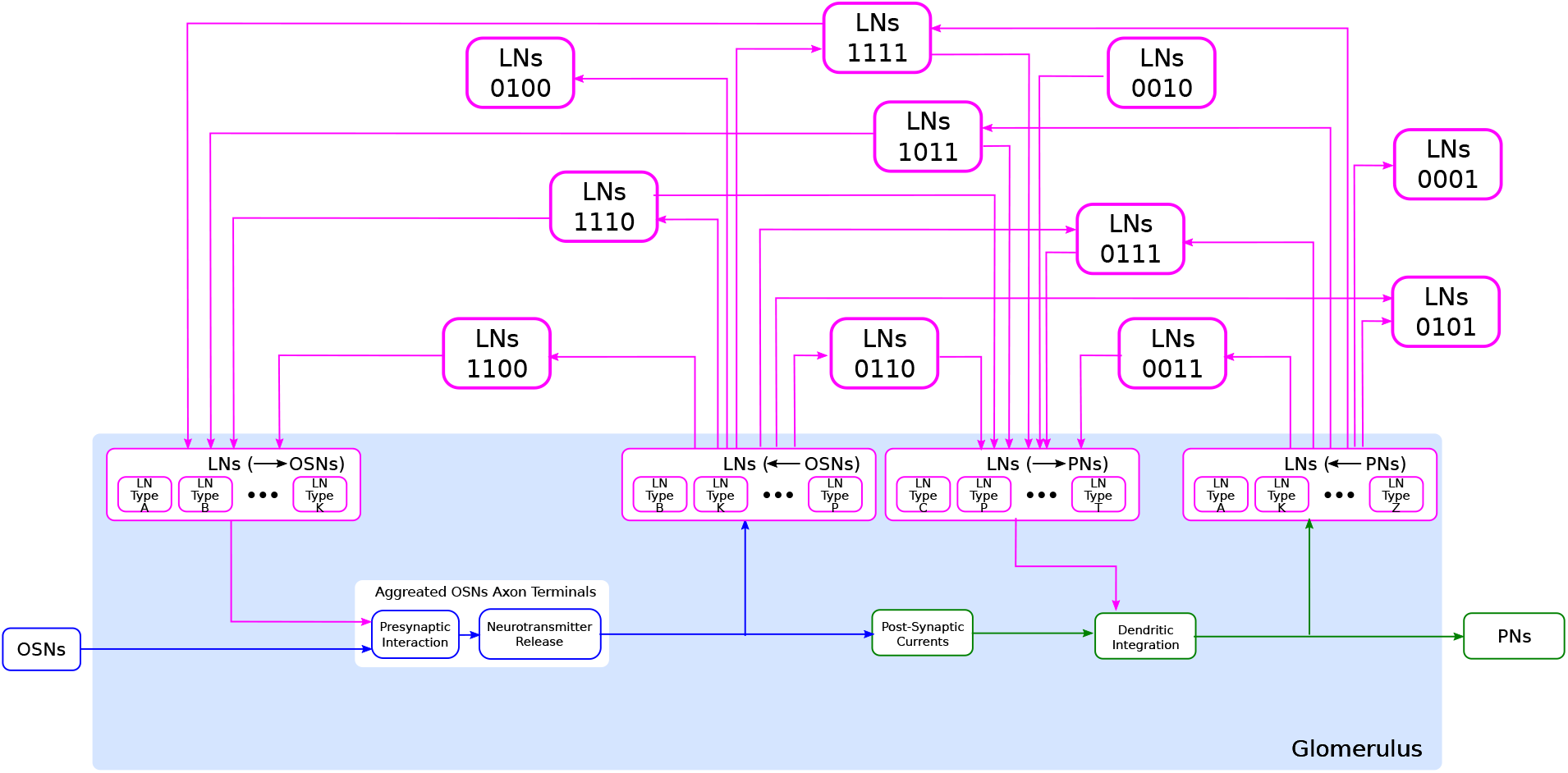
Schematic diagram of a single glomerulus circuit. (bottom) The blocks within the glomerulus represent the feedforward circuits. (top) LNs with different interaction connectivity patterns. Interaction connectivity patterns that have less than 50 occurrences in the Hemibrain dataset are omitted (see also Table 1).

Since the OSN axon terminals and PN dendrites are all inside the respective glomerulus they project into, their interaction with the LNs must also occur within the same glomerulus. Therefore, in the circuit diagram of the glomerulus in Figure 8, we included 4 types of interactions between OSNs and LNs and between PNs and LNs. First, LN outputs interact presynaptically with OSN axon terminals and modulate their neurotransmitter release upon incoming spikes [24]. Second, LNs receive inputs directly from OSN axon terminals. Third, LNs provide inputs to PNs. Finally, LNs receive inputs from PN dendrites. Within the glomerulus, however, we do not specify the exact LNs that carry out these interactions. Rather, we define 4 ports (see magenta blocks in Figure 8: i) LNs (→OSNs), ii) LNs (←OSNs), iii) LNs (→PNs), and iv) LNs (←PNs), corresponding to, respectively, the 4 types of interactions mentioned above. The connections from/to the specific LNs will be defined through these ports. All the LNs that connect to each port carry out the specific interaction connectivity pattern within the glomerulus.

#### Modeling LN Feedback Circuits

Each LN has its own connectivity pattern onto the ports of a glomerulus. Given the 4 ports in each glomerulus, there are 15 different possible connection patterns that an LN can interact with within a given glomerulus. We use a 4 digit binary code to represent this interaction, according to the left-right order of the ports in Figure 8. For example, if an LN receives inputs from OSNs and provides feedback to the same OSNs, but has no interaction with PNs, then we call the interaction connectivity pattern of this LN with the glomerulus of type “1100”. Figure 8 shows 11 of the 15 types of most common LN connections with a glomerulus. Note that a single LN can have different types of interaction connectivity patterns with different glomeruli. Therefore, this code does not define LN types but rather the interaction connectivity pattern within a given glomerulus.

Inspecting all 226 LNs that innervate the right AL in the hemibrain dataset [8], we list in the 2nd column of Table 1 the number of instances each interaction connectivity pattern occurs across 51 olfactory glomeruli. For the DM4 and DL5 glomeruli, the number of occurrences of each interaction connectivity pattern is listed in the 3rd and 4th column, respectively.

**Table 1:** Number of occurrences of interaction connectivity patterns in the AL. 2nd column: Number of occurrences for all LN innervations in all 51 olfactory glomeruli, where each LN innervation in a glomerulus counts as 1 occurrence. 3rd column: Number of occurrences in the DM4 glomerulus (*i.e.*, the number of LNs that have an interaction connectivity pattern in DM4). 4th column: Number of occurrences in the DL5 glomerulus (*i.e.*, the number of LNs that have an interaction connectivity pattern in DL5).

| Interaction<br>Connectivity<br>Pattern | # of<br>Occurrences<br>(all glomeruli) | # of<br>Occurrences<br>(DM4) | # of<br>Occurrences<br>(DL5) |
| --- | --- | --- | --- |
| 1111 | 808 | 21 | 26 |
| 0011 | 725 | 13 | 19 |
| 0111 | 515 | 14 | 8 |
| 0100 | 318 | 8 | 4 |
| 0010 | 263 | 1 | 4 |
| 1110 | 239 | 6 | 7 |
| 0001 | 221 | 4 | 0 |
| 0110 | 131 | 7 | 2 |
| 0101 | 106 | 4 | 1 |
| 1011 | 69 | 0 | 0 |
| 1100 | 60 | 0 | 0 |
| 1101 | 30 | 1 | 0 |
| 1000 | 18 | 1 | 0 |
| 1010 | 16 | 0 | 0 |
| 1001 | 3 | 0 | 0 |

For a single glomerulus, an LN is considered to form a self-feedback loop if the interaction connectivity pattern is 11xx, xx11 or 1xx1. They account for 8 out of 15 interaction connectivity patterns. The rest of the connectivity patterns are involved in cross-feedback loops from/to other glomeruli. In the case when an odorant only activates a single type of OR and hence excites only 1 group of OSNs, the self-feedback loops shape the response of the PNs. Although this case rarely occurs naturally, it is possible to experimentally validate our model of a single glomerulus with self-feedback by optogenetically activating only a single group of OSNs expressing the same OR [25, 26].

#### Abstraction of Glomerular Feedback Circuit Motifs

While the above model of LN feedback loops is based on connectome data, it is difficult to study individual interaction connectivity patterns even for a single glomerulus. This is largely due to the massive number of LNs and the wide variety of interaction connectivity patterns associated with each glomerulus.

Here, we abstract all LN interaction connectivity patterns into three feedback motifs as depicted in Figure 9.

**Figure 9:**
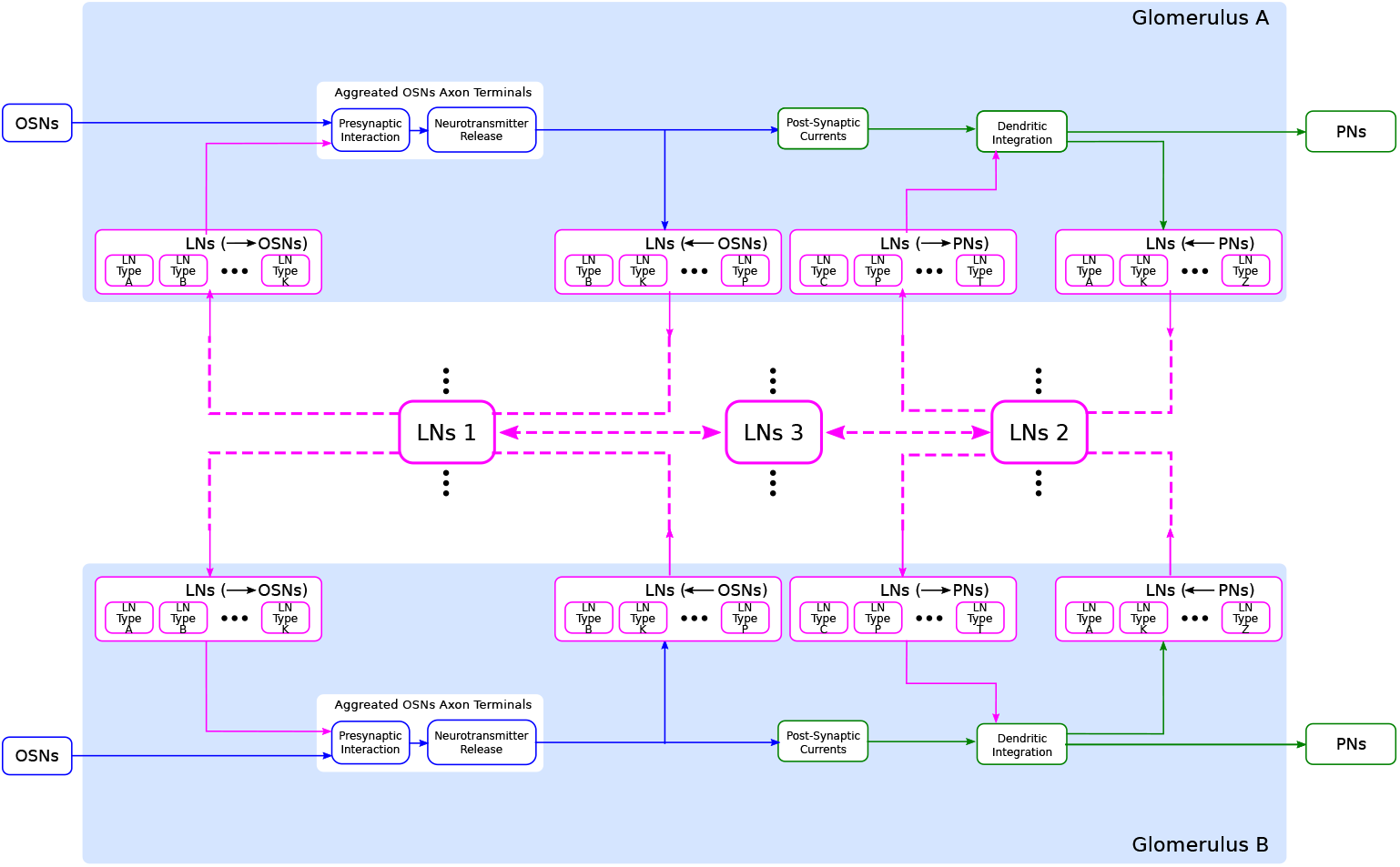
An abstraction of glomerular feedback circuits with 3 feedback motifs. Feedback motif 1 (LNs 1) forms the feedback loops on OSNs. Feedback motif 2 (LNs 2) forms the feedback loops on PNs. Feedback motif 3 (LNs 3) forms the feedback loop between the other two loops.

The first feedback motif models the interaction connectivity patterns of LNs with OSNs. The LNs of feedback motif 1 receive inputs from OSN axon terminals and feed back into the same OSN axon terminals.

The second feedback motif models the interaction connectivity patterns of LNs with PNs. The LNs of feedback motif 2 receive input only from PNs and feed back into the same PNs.

The separation of the two loops above allows us to address the individual contribution of the feedback motifs, respectively, in controlling the inputs into the glomeruli and in reshaping the glomerular output onto the LNs.

Finally, the third feedback motif models the interaction between the above two loops through additional LN-to-LN connectivity patterns. The LNs of feedback motif 3 do not receive or feedback to either OSNs or PNs. Rather, they connect only with LNs of feedback motif 1 and 2. Such a feedback motif enables the glomerular outputs and the glomerular inputs to influence the odor signal processing of other glomeruli.

#### Modeling the Antennal Lobe Feedback Circuits

With models of a single glomerulus and LN feedback motifs, we can generalize these feedback circuits to the entire AL. In particular, the morphological LN types described in Section 2.1 provide a blueprint for connecting multiple glomeruli via the LNs. Figure 10(A) depicts one such LN that innervates more than 20 glomeruli.

**Figure 10:**
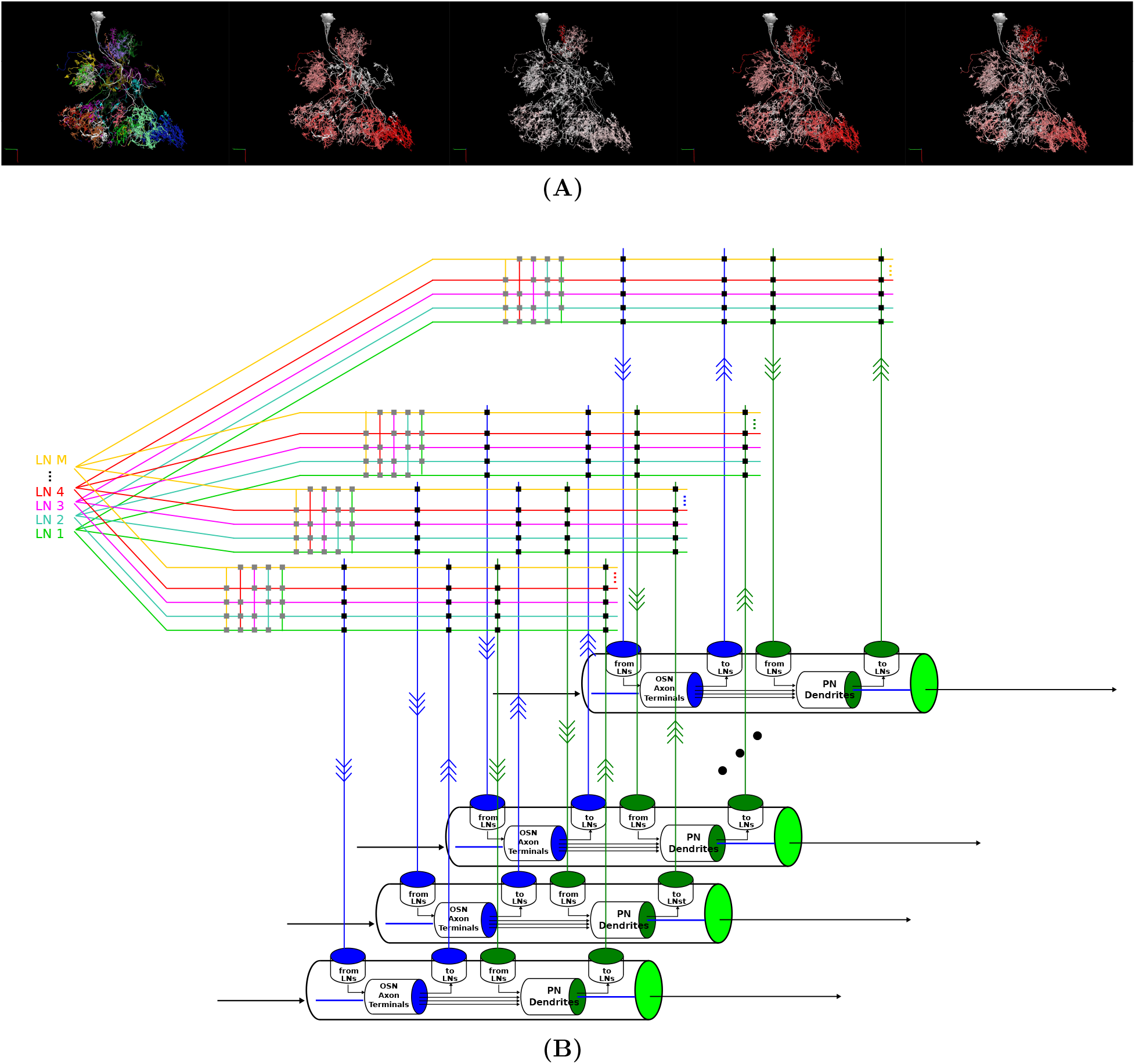
A circuit diagram modeling the entire AL feedback circuit. **(A)** Morphology of an LN and its connectivity to the glomeruli it innervates. From left to right: 1) LN innervation of glomeruli. Each color indicates the glomerulus it arborizes. 2) Number of synaptic inputs from OSNs. The arbors of the LN in each glomerulus is colored in red with saturation proportional to the number of synapses, *i.e.*, redder indicates higher number of synapses. 3) Number of synaptic outputs to OSNs. 4) Number of synaptic outputs to uPNs. 5) Number of synaptic inputs from uPNs. **(B)** Schematic diagram of the overall feedback circuit in the AL.

To model the entire AL feedback circuit, we define glomeruli as parallel channels [27] each exposing 4 ports that were depicted in Figure 8. The ports and the LNs form a crossbar as depicted in Figure 10(B), where the crossbar “intersections” are marked with black dots. LNs also form a second crossbar associated with each glomerulus, depicted in Figure 10(B), where the crossbar “intersections” are marked with grey dots. The exact connectivity pattern can be determined for specific connectomic datasets.

## 4 Interactive Exploration of the Functional Logic of Feedback Circuits

In this section, we present an approach for exploring the functional logic of feedback circuits in the fruit fly brain. This pertains to the third column of Figure 2. Specifically, we present here the interactive exploration of the AL following the previous section where the AL feedback circuits have been explored (Section 2) and modeled (Section 3).

We describe a library for instantiating antennal lobe feedback circuit motifs from connectome data with customizable parameters of neurons and synapses. We then demonstrate the use of this circuit library in exploring the I/O of a single glomerulus as well as two interconnected glomeruli. We also provide an outline of scaling the methodology presented here to the entire AL circuit.

### 4.1 Circuit Library for Exploring the Functional Logic of the Massive Number of Feedback Loops in the Antennal Lobe

We introduce here a circuit library, called FeedbackCircuits, for exploring the functional logic of the massive number of feedback loops (motifs) in the fruit fly brain. While the library is generic and can applied to any local processing unit of the fly brain, we highlight here its capabilities in constructing and exploring the AL feedback circuit models described in Section 3.

First, the FeedbackCircuits Library provides tools for interactively visualizing and exploring the feedback loops in the AL circuit operational on the FlyBrainLab computing platform [16] (see Materials and Methods for details).

Second, the FeedbackCircuits Library enables users to instantiate an executable circuit of the feedback circuit model in two ways. An executable circuit can be instantiated according to a connectome dataset. For example, any circuit explored via NeuroNLP++ can be loaded into an executable circuit directly. It can also be instantiated according to the abstraction of feedback motifs defined in Section 3.3 (see also Materials and Methods).

While the connectivity pattern of neurons can be extracted from connectome datasets, users can also define higher level objects, such as glomeruli of the AL (see also Materials and Methods). Within a chosen object, the characteristics of the executable models, such as the dynamics of neurons of different cell-types, are specified by the user. For example, users can specify all OSN to PN connections to exhibit commonly-used synaptic dynamics. Every instance of such synapses, residing in the connectome dataset, will be automatically assigned the specified dynamics. Similarly, all LN to OSN connections can be specified to act presynaptically on OSN axon terminals [28].

Finally, LNs of different types, LNs with different connectivity patterns or different LN motifs can be flexibly ablated in the FeedbackCircuit Library and their individual and combined effect on the AL outputs evaluated.

The FeedbackCircuits Library provides easy-to-customize loader and visualization functions to explore the I/O behavior of the antennal lobe circuit. This process can be repurposed for a wide variety of neuropils, including the mushroom body and the lateral horn of the early olfactory system.

To demonstrate the capabilities of the FeedbackCircuits Library, we explore below the I/O of the glomerular circuit abstraction described in Section 3.3. We start with a one-glomerulus scenario, then extend our findings to a two-glomeruli scenario and finally show the ease with which the contribution of different neurons or feedback loops to the overall circuit function can be interrogated.

### 4.2 Exploration of the Functional Logic of Feedback Circuits of a Single Glomerulus in Isolation

In Figure 11, we evaluate the I/O behavior of the DM4 and DL5 glomeruli for different combinations of LN motifs. We outline how the presence of different feedback motifs can jointly or individually alter the PN output of the glomeruli. The number of OSNs and PNs, as well as the connectivity between these two types of neurons are configured according to the Hemibrain connectome dataset. In Figure 11(A, B), we show the correspondence of these LN motif exemplars to real LNs in the Hemibrain dataset. LNs depicted in Figure 11(A) form feedback loops with OSNs and PNs in the DM4 glomerulus. Those shown in Figure 11(B) form feedback loops with the OSNs and PNs in DL5 glomerulus.

**Figure 11:**
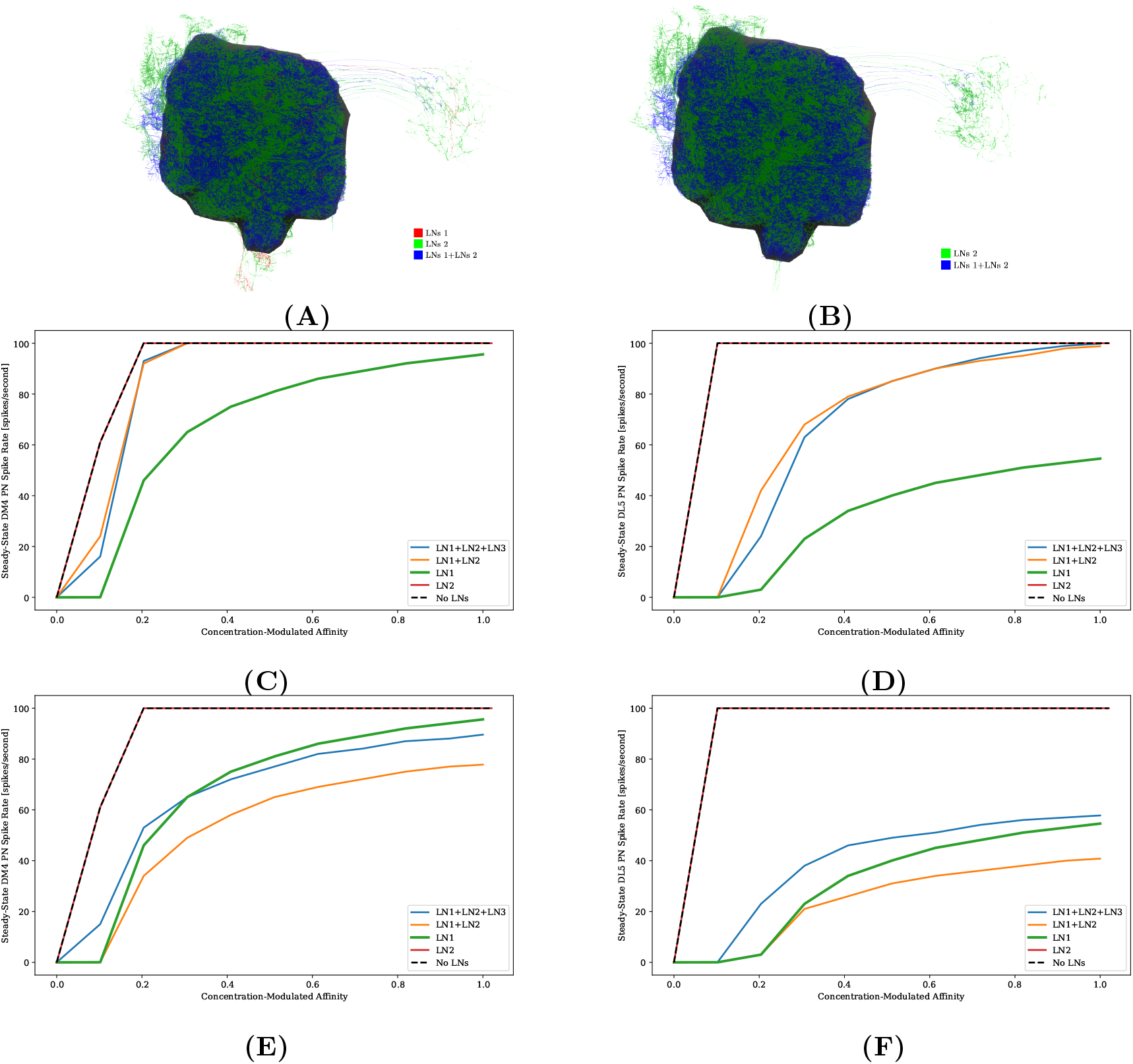
Characterization of PN responses of the single (isolated) DM4 and DL5 glomerular circuits. **(A)** LNs that form LN1 (red, 2 LNs), LN2 (green, 54 LNs) feedback loops with the DM4 glomerulus and that exhibit both both LN1- and LN2-type feedback characteristics (blue, 18 LNs). LN3-type feedback loops are not shown. **(B)** LNs that form LN2 (green, 48 LNs) feedback loops with the DL5 glomerulus and that exhibit both LN1- and LN2-type feedback characteristics (blue, 18 LNs). LN3-type feedback loops are not shown. **(C)** DM4 PN steady-state firing rate across different constant odorant concentration levels with DM4 executed in isolation. (dashed black) No LN is present. (green) Only motif 1 LNs are present. (red) Only motif 2 LNs are present. (orange) Motif 1 and 2 LNs are present. (blue) Motif 1, 2 and 3 LNs are present. **(D)** DL5 PN steady-state firing rate across different constant odorant concentration levels with DL5 executed in isolation. Colors are the same as in (A). Motif 2 LNs in both (A) and (B) are assumed to be excitatory. **(E)** DM4 PN steady-state firing rate with DM4 executed in isolation and Motifs 2 LNs assumed to be inhibitory. Colors are the same as in (A). **(F)** DL5 PN steady-state firing rate when DL5 is executed in isolation and Motifs 2 LNs assumed to be inhibitory. Colors are the same as in (A).

In Figure 11(C, E), we evaluate and compare the DM4 glomerulus response due to different combinations of LN feedback motifs. Following the motifs we explored in Figure 9, we added to the single DM4 glomerular circuit composed of the OSNs and PNs three LN motifs: LN1 (feedback motif 1), LN2 (feedback motif 2) and LN3 (feedback motif 3) (see also Materials and Methods). Results shown in Figure 11(C) were obtained assuming that the LN2 is excitatory, while those in Figure 11(E) assuming that the LN2 is inhibitory.

For constant odorant concentration waveforms, the steady-state firing rates of the corresponding PNs are displayed in Figure 11(C) (see also Materials and Methods). For the range of binding affinities evaluated, if the glomerulus is configured without any feedback loops it is driven immediately to saturation (Figure 11(C) dashed black curve). Note that the curve graphically displaying the firing rates is clipped to a maximum of 100 spike/s. The addition of feedback motif 1 LNs that presynaptically inhibit OSNs quickly, results in a sigmoidal PN spiking rate for the tested range of odorant concentration values (Figure 11(C) green curve). LNs exhibiting this feedback motif are observed in all glomeruli, suggesting an important role in regulating the glomerular output by this feedback motif. We also note that the addition of feedback motif 2 alone, either excitatory or inhibitory, does not directly contribute to regulating the PN response, and results in a saturation level similar to the circuit without feedback (Figure 11(C,E) red curves).

We evaluate next the effect of the combination of feedback motifs. The orange curves in Figure 11(C) and (E) depict the responses of DM4 PNs when, respectively, excitatory and inhibitory LN2 is added to the circuit with the LN1 feedback motif already present. Finally, we add LN3 on top of the previous setup and stimulate this new neuron externally with a 20nA current source to be able to test how activation of this neuron affects the responses of the DM4 PN; this results in regular spiking in LN3 and suppression of both LN1 and LN2; but the suppression of LN2 causes a larger effect and thus a net decrease in the PN spiking rate. These simple explorations demonstrate the effect of different feedback loop configurations might have on the responses of the one-glomerulus circuit.

Evaluations with the same set of configurations were performed on the feedback circuit of the DL5 glomerulus (Figure 11(D) where LN2 is excitatory and Figure 11(F) where LN2 is inhibitory), in which we observe similar contributions from different configurations of the feedback motifs as in the case of the DM4 glomerulus. Results for DM4 and DL5 if LN2 is inhibitory are shown in Figure 11(E) and Figure 11(F), respectively. As expected, in this scenario, ablation of LN2 causes a higher spike rate of the PN. An indirect suppression of LN2 through stimulation of LN3 similarly raises the spike rate of the PN.

The comparison of the PN outputs using different feedback motifs (Figure 11(C-F)) shows that i) the LN1 feedback motif is essential for the circuit to be stable under a large range of concentration amplitude values, ii) the addition of the LN2 feedback motif amplifies the steady-state PN spike rate response given the presence of LN1 feedback motif, and LN3 feedback motif controls the contribution of LN1 and LN2 on the circuit.

### 4.3 Exploration of the Functional Logic of Feedback Circuits of Two Interconnected Glomeruli

In Figure 12, we evaluate a circuit consisting of interconnected DM4 and DL5 glomeruli. The circuit diagram of the interconnected DM4 and DL5 glomeruli according to the Hemibrain dataset is shown in Figure 12(A). For simplicity in evaluation, we again used the abstraction presented in Section 3.3. The number of OSNs and PNs in these two glomeruli, and the number of synapses between the two types of neurons are configured according to the Hemibrain dataset. We then add five LNs: 1 LN1 each connects only to DM4 and DL5, 1 LN2 each connects only to DM4 and DL5, and 1 LN3 connected to all other LNs in both directions, following the motifs we explored in Figure 9 (see also Materials and Methods).

**Figure 12:**
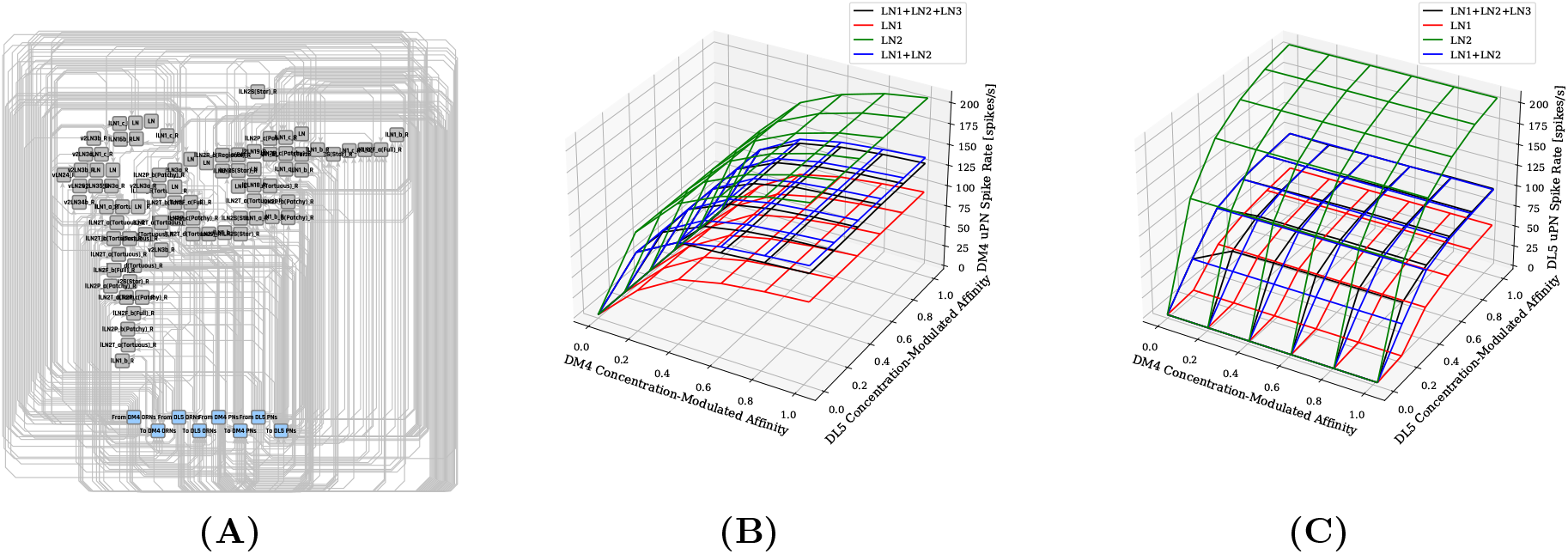
Characterization of DM4 and DL5 PN responses of a circuit consisting of interconnected DM4 and DL5 glomeruli. **(A)** A circuit diagram of the interconnected DM4 and DL5 glomeruli. Towards the bottom, the blue nodes correspond to ports to and from DM4/DL5 OSNs/PNs. Each gray node corresponds to a specific LN from the Hemibrain dataset. Only connectivity between the gray nodes and the blue nodes are shown in the diagram. **(B)** DM4 PN steady-state firing rate and **(C)** DL5 PN steady-state firing rate as glomeruli are subject to different odorant concentration values. For both (B) and (C): (red) Only motif 1 LNs are present. (green) Only motif 2 LNs are present. (blue) Motif 1 and 2 LNs are present. (black) Motif 1, 2 and 3 LNs are present.

In Figure 12(B), we show the average spike rate of the DM4 PN as a function of the concentration-modulated affinities (see Materials and Methods for an interpretation of the inputs). In Figure 12(C), we show the average spike rate of the DL5 PN as a function of the concentration-modulated affinities. Mirroring Figure 11, we consider combinations of different subsets of LNs. Similar to Figure 11, we find that the addition of LN1 alone produces the lowest spiking rate (Figure 12(B,C) red mesh). Similarly, the addition of LN2 alone produces the highest spiking rate due to lack of presynaptic inhibition from LN1 (Figure 12(B,C) green mesh). When we added feedback motif 2 in addition to feedback motif 1, the excitatory nature of the LN2 loop resulted in the spike rate increasing by a small amount when compared with feedback motif 1 alone (Figure 12(B,C) blue mesh). Finally, we added LN3 to the previous setup; we see that the spike rate for a given glomerulus increases with the affinity of the receptors expressed by the OSNs projecting to that glomerulus. Note that, an increase in the odorant input amplitude of the OSNs projecting to the other glomerulus also affects the spike rate through LN3 (Figure 12(B,C) black mesh).

The abstraction we considered here can be extended to the entire AL through the Feedback-Circuits Library that we provide. The exact interconnect of different LN feedback motifs with each glomerulus can be extracted from the connectivity of LNs in the Hemibrain connectome. Cross-glomerular effects can be tabulated given a computational hypothesis about the circuit.

## 5 Discussion

### A Programmable Ontology Encompassing the Functional Logic of the Brain

Existing neuroscience-centered ontologies, including those of the fruit fly [15], the rat/mouse [29, 30, 31] and the human [32] brain, mainly focus on neuroanatomical structures, hierarchies and nomenclature. Description of entities or relations that have functional significance is rare and is kept at behavioral or cognitive level [33].

While describing the structure of the brain is certainly a first step in the quest of understanding brain function, it is far from being sufficient. Thus, an ontology of the brain can not end with the description of anatomical data. Rather, the anatomical entities and relations have to be augmented with insights characterizing the functional logic of brain circuits.

In this paper, we presented a programmable ontology that expands the scope of the current ontology of *Drosophila* brain anatomy [15, 16] to encompass the functional logic of the fly brain. The programmable ontology provides a language not only for modeling circuit motifs but also for programmatically exploring their functional logic. To achieve this goal, we tightly integrated the programmable ontology with the workflow of the interactive FlyBrainLab computing platform [16]. In effect, the programmable ontology, embedded into the FlyBrainLab, has grown into a programming environment operating with access to a plethora of datasets, containing models of sensory space, the connectome/synaptome including cell types/feedback loops and neuronal/synaptic dynamics. The programmable ontology has the “built in” capability for evaluating the functional logic of brain circuits and for comparing their behavior with the biological counterparts.

To provide a language for defining functional circuit motifs anchored onto biological datasets and the worldwide literature, we developed the NeuroNLP++ web application that supports free-form English query searches of ontological entities and references to these in the published literature worldwide. NeuroNLP++ enables circuits to be composed using connectomic/synaptomic data in support of the evaluation of their function *in silico*. To bridge the gap between the existing *Drosophila* Anatomy Ontology dataset and the Hemibrain connectome morphology dataset, we associated with each ontological entity the corresponding neurons in the morphology dataset. The DrosoBOT Engine, in conjunction with the rule-based NLP engine, represents a first step towards providing a unified and integrated view of connectomic/synaptomic datasets and of the fruit fly brain literature worldwide.

In our programmable ontology the modeling of the space of sensory stimuli is explicitly included. We note that, e.g., the space of odorants has not been discussed in formal ontologies of the fly brain anatomy, although it plays a key role in defining, characterizing and evaluating the functional logic of brain circuits. Here, the odorant space is modeled by a 3D tensor trio that describes the interaction between odorants and olfactory receptors, rather than by the (largely intractable) detailed/precise chemical structure of the odorants. Defining odorants and odorant mixtures as well as their interactions with olfactory receptors is an important step of this program.

By augmenting the ontology with the space of odorant objects and by providing an English query web pipeline for exploring structural features of the architecture of the early olfactory brain circuits, we are now in a position to evaluate the functional logic of these circuits in their full generality. The program of research presented here was, due to space limitations, set the stage to modeling and evaluation of the early olfactory system. Clearly, an extension to the other sensory modalities is in order. In particular, the early vision system [22] and the central complex [34, 16] are our next candidates.

### Construction of Circuit Motifs with the FeedbackCircuits Library

Detailed connectomic datasets, such as the Hemibrain dataset, reveal a massive number of nested feedback loops among different cell-types. Dissecting the role of these feedback circuits is key to the understanding the model of computation underlying the Local Processing Units (LPUs) of the fruit fly brain. The methodology underlying the FeedbackCircuits Library we advanced here has wide reaching implications for studying the massive feedback loops that dominate all regions/neuropils the fruit fly brain.

The FeedbackCircuits Library brings together the available *Drosophila* connectomic, synaptomic and cell-type data, with tools for 1) querying connectome datasets that automatically find and incorporate feedback pathways, 2) generating interactive circuit diagrams of the feedback circuits, 3) automatic derivation of executable models based on feedback circuit abstractions anchored on actual connectomic data, 4) arbitrary manipulation (and/or ablation) of feedback circuits on the interactive circuit diagram for execution, and 5) systematic characterization and comparison of the effect of different feedback loops on the I/O relationship of arbitrary brain circuits.

We have demonstrated the capabilities of the FeedbackCircuits Library using circuits of the DM4 and DL5 glomeruli of the *Drosophila* antennal lobe constructed, based on the Hemibrain dataset, either individually in isolation or jointly interconnected. We have demonstrated the methodology to construct and explore these feedback circuits to characterize the contribution of individual feedback motifs as well as their compositions. Constructing the entire AL feedback circuit is also in sight. The methodology advanced here paves the way to investigate similar feedback circuits in other regions of the fly brain, such as the medulla, the MB, etc. The studies of the functional logic of sensory processing in these neuropils are currently limited to the feedforward pathways [35, 36, 37], although these circuits exhibit strong feedback components [38, 39, 16].

## 6 Materials and Methods

In this section, we present the details of the methodology for the exploration of the morphology of massive feedback circuits, modeling odor signal processing in the early olfactory system, and the interactive exploration of the antennal lobe as a network of glomeruli.

For creating the tools underlying the programmable ontology we used extensive capabilities to query datasets and build executable circuits, query the antennal lobe circuitry using these as well as customized tools, constructing and evaluating the feedback circuits with the FeedbackCircuits Library, and mapping glomeruli and their compositions into executable circuits.

### 6.1 Exploring the Morphology of Massive Feedback Circuits

#### NeuroNLP++

NeuroNLP is a web application that supports the exploration of fruit fly brain datasets with rule-based English queries [17, 16]. To enhance the user experience when asking questions that are well beyond the current capabilities of NeuroNLP, we devised the NeuronNLP++ brain explorer (https://plusplus.neuronlp.fruitflybrain.org). Figure 13 depicts the software architecture of the NeuroNLP++ application. In addition to the backend servers supporting the NeuroNLP web application (NeuroArch Server and NeuroNLP Server with rule-based NLP Engine), NeuroNLP++ is supported by the DrosoBOT Engine (see below), i.e., an additional backend of the NeuroNLP Server.

**Figure 13:**
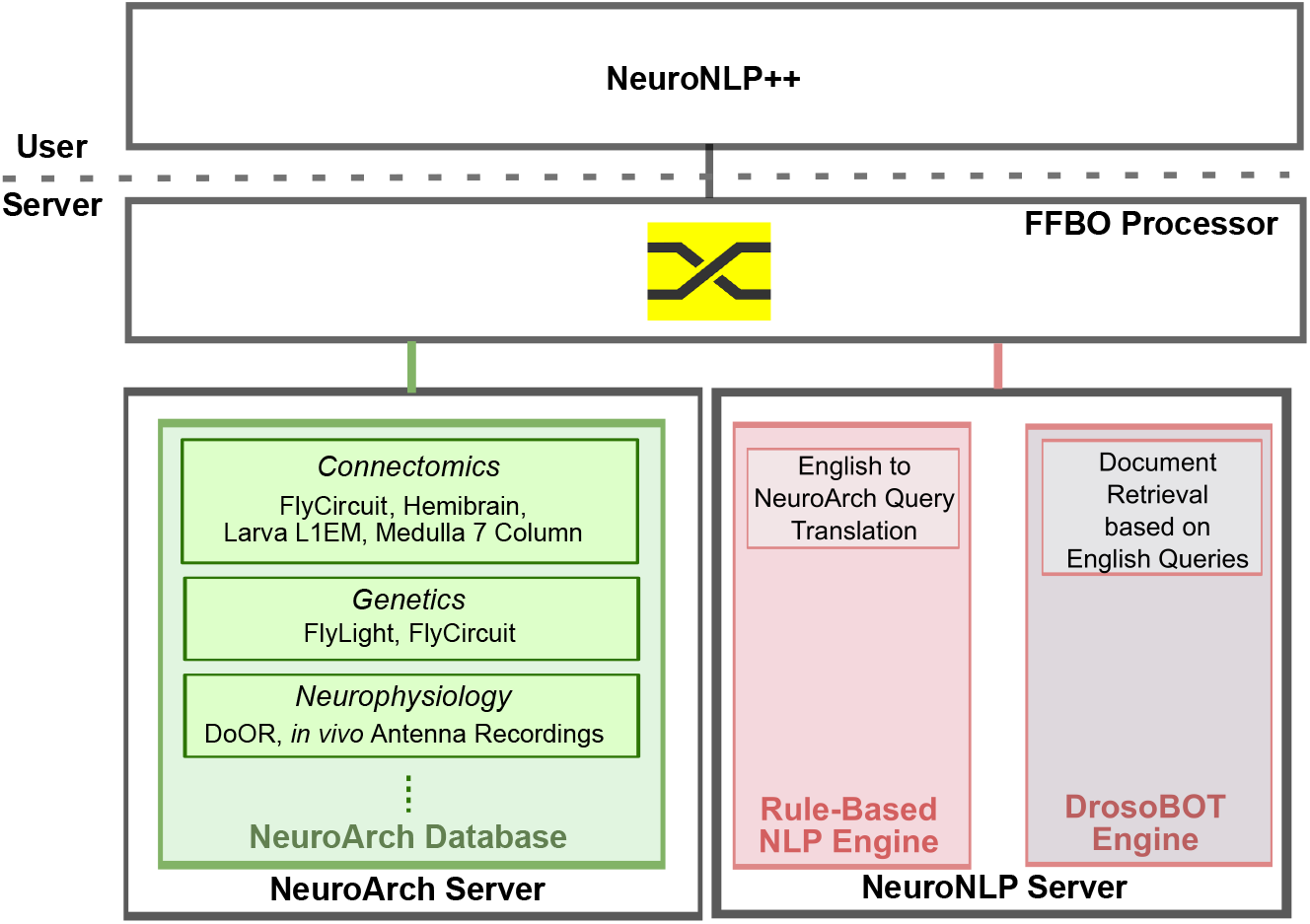
Software architecture of the NeuroNLP++ web application. The frontend application communicates with three backend servers: a NeuroArch Server that hosts a NeuroArch Database of fruit fly brain data, a NeuroNLP Server that runs the rule-based NLP engine, and, an additional NeuroNLP Server running the DrosoBOT Engine interpreting for free-form English queries.

When the query is a free-form English question that cannot be interpreted by the rule-based NLP engine, it is sent to DrosoBOT. DrosoBOT responds with a list of ontological entities that are most pertinent. Each entry in the list includes the name of the cell-type, a link to the *Drosophila* Anatomy Ontology containing references to the entities in question [15], and a description of the cell-type as well as relevant entries to the worldwide literature (see also Figure 3(A)). In addition, it includes buttons to add, pin and unpin the neurons in the 3D visualization workspace.

In addition to using DrosoBOT to resolve free-form English queries, NeuroNLP++ includes a “graph view” functionality that allows users to visualize the graph representing the connectivity of neurons in their workspace at single-cell or cell-type level. Once the graph view button is pressed, NeuroNLP++ retrieves the connectivity of all the neurons in the workspace, with an additional capability to filter out the connections that have less than *N* synapses, where *N* ≥ 0. A graph is then plotted in the workspace using the sigma.js library.

For a single-cell level graph, each node represents a single neuron in the visualization workspace, and the edge between two nodes represents the number of synapses between the two corresponding neurons. For a cell-type level graph, each node represents a cell-type that may include multiple neurons of the same cell-type in the visualization workspace. The number of synapses from all the neurons in one node (cell-type) to all the neurons in another node (cell-type) are accumulated to form an edge between the two nodes. In addition, the graph in the “Graph View” is interactive. Hovering the mouse on a node highlights the corresponding neuron or all neurons of the corresponding cell-type in the 3D visualization. The graph can be further rearranged by hand.

#### The DrosoBOT Engine

DrosoBOT is a natural language processing engine that 1) parses free-form English queries pertaining to entities available in an ontological dataset [15], and 2) provides morphological data from a connectome dataset [8] already associated with each ontological entity.

Given a free-form English query, DrosoBOT first uses DPR to retrieve relevant passages in the query as context candidates, and then uses PubMedBERT fine-tuned on the Stanford Question Answering Dataset to find possible answers to questions pertaining to a collection of *Drosophila*-specific ontology terms and their descriptions. Here DPR is the dense passage retriever trained on the Natural Questions dataset [40] that uses real anonymized queries issued to Google and annotated answers from the top 5 Wikipedia articles. PubMedBERT is the Bidirectional Encoder Representations from Transformers (BERT) [41] model with biomedical domain-specific pre-training [42] from abstracts on PubMed.

In addition, for specific cell-types, DrosoBOT implements a modular lexical search subsystem that uses domain knowledge to improve search results when specific keywords of cell-types are asked. We make use of this system to improve the search results for the antennal lobe, which requires biological nomenclatures such as “DM4” to be detected not as typos but as important structures.

To bridge the gap between existing ontology and connectome datasets, we associated with each ontological entity the corresponding neurons in the Hemibrain dataset based on the names of the entities and their synonyms after searching through all possible matches in the *Drosophila* Anatomy Ontology (DAO) dataset [15]. We then created a graph with nodes consisting of both names of entities in the DAO and names of neurons. An edge is created between two nodes with a matching term. After finding the ontological term relevant to the English query from the first step, we then retrieved the names of the neurons that are the graph neighbors of the ontological entity, and finally retrieved the neurons from the database.

For the AL, starting with the terms for cell-types and abstractions in [15] and expanding these to include references to all cell-types so that all common synonyms are accounted for (for example, PNs, OSNs, glomeruli and LNs), we facilitated the specification of antennal lobe circuits through queries. Here we provided the capability to add relevant groups of neurons such as new glomeruli and local neurons in only a few searches and button presses. We also added the names of all glomeruli as special ”keywords” whose association with the antennal lobe is automatically detected if present in a search query. This hybrid approach with rule-based detection of special keywords and neural searches allows for terms relating to antennal lobe to be retrieved whilst keeping the search engine open for more documents and keywords. The latter can readily be edited by DrosoBOT developers.

#### Defining Feedback Circuits in the Antennal Lobe

In order to compute anatomical correspondences to LN1- and LN2-type feedback loops in each AL glomerulus, we first extracted all OSNs that project to a glomerulus and all the PNs that innervate that glomerulus. We consider that an LN forms an LN1-type feedback loop with OSNs if it receives a total of more than 5 synapses from all the OSNs and provides a total of more than 5 synapses to all the OSNs, and it has less than 5 synapses with all the PNs. Similarly, we consider an LN forms an LN2-type feedback loop with PNs if it receives from and provides to all the PNs a total of more than 5 synapses and has less than 5 synapses with all the OSNs. An LN is considered to exhibit both LN1- and LN2-type feedback characteristics if it receives from and provides to all the OSNs a total of more than 5 synapses and if it receives from and provides to all the OSNs a total of more than 5 synapses. The LNs identified with each type of feedback loop in a glomerulus are then associated with an ontological entity that is accessible by DrosoBOT for document retrieval.

### 6.2 Modeling the Space of Odorants and Early Olfactory Processing

#### Modeling the Space of Odorants

To construct odorant objects as elements of the space of odorants, we fitted, by following [21], the affinity tensor **b***/***d** with entries [**b**]_*ron*_/[**d**]_*ron*_, for all *r* = 1, 2*, ..., R*, *o* = 1, 2*, ..., O* and *n* = 1, 2*, ..., N*. to match the steady-state spike rate in response to a constant waveform of 110 different odorants [43]. Detailed data can be found in the olfactory transduction circuit library, OlfTransCircuit see https://github.com/FlyBrainLab/OlfTrans [16].

#### Modeling and Constructing the Massive Feedback Circuits of the AL

To identify the interaction connectivity patterns of the LNs leading to the circuit diagram in Figure 8, we inspected all 311 LNs in the Hemibrain dataset [8]. Of these, 296 LNs have more than 10 synapses in the right hemisphere AL. We only considered a synapse if both its presynaptic and postsynaptic sites are identified at a confidence level higher than 70% in the Hemibrain dataset. For each of these LNs, we counted the number of synapses they make, presynaptically and postsynaptically, with partner OSNs as well as PNs in each glomerulus. If the total number of synapses within a glomerulus is less than 5, we deem the connectivity pattern to be 0000, *i.e.*, no connection pattern. The first digit of the 4-digit binary code is 1 if the number of LN to OSNs synapses is larger than 5. Similarly, the second digit is 1 if the number of synapses the LN receives from OSNs is larger than 5. The third digit is 1 if the number of LN to PNs synapses is larger than 5. Similarly, the fourth digit is 1 if the number of synapses the LN receives from PNs is larger than 5.

Construction of the circuit diagram of the entire AL in Figure 10 comes in three steps. First, OSNs and PNs and their ports from/to LNs are constructed for each glomerulus according to Figure 8. ome of the glomeruli are depicted as cylinders at the bottom of Figure 10(B). Second, for each glomerulus, we also configure the interaction connectivity patterns of LNs, as described above. This forms the local crossbar between all LNs and the ports of a glomerulus and is depicted, e.g., on the top right of Figure 10(B), as 4 vertical dotted lines. Finally, connecting all these local crossbars from all glomeruli based on the innervation pattern of each LN, we obtain a hierarchical crossbar between LNs and the ports of glomeruli. The hierarchical crossbar provides the flexibility to configure the routing of interconnections across glomeruli, either by using the connectivity patterns of LNs extracted from from connectome data, or by any variations/ablations thereof for testing and evaluating the functional logic of the AL circuit.

### 6.3 Exploring the Function of the AL as a Network of Glomeruli

#### Interactively Exploring Circuit Diagrams with the FeedbackCircuits Library

The FeedbackCircuits Library is developed in Python and designed to be integrated into the FlyBrainLab ecosystem for constructing feedback circuits and exploring their function.

The FeedbackCircuits Library provides tools for interactively visualizing and exploring the feedback loops of the AL circuit operational on the FlyBrainLab computing platform [16]. Figure 14(A) shows a typical FlyBrainLab user interface. On the top left is the NeuroNLP 3D visualization window for displaying the morphology of neurons. At the bottom left a Jupyter notebook for code execution is displayed. On the right an automatically generated diagram of the DM4 glomerulus circuit is shown. This circuit diagram is a schematic of the glomerulus model shown in Figure 8 and is based upon the Hemibrain connectome dataset [8]. The LNs that have different interaction connectivity patterns with the DM4 glomerulus are grouped into blocks. The circuit diagram allows users to inspect the morphology of neurons in the NeuroNLP window by clicking on the ones displayed. For example, in Figure 14(B, right), we zoomed into the LNs that exhibit the connectivity pattern 0101. By clicking on the neuron whose Hemibrain ID is 1702323388, the LN in green is highlighted in Figure 14(B, top left). Hovering also highlights all connected neurons, such as the one with Hemibrain ID 1825789179 in Figure 14(B, right). The interactive circuit diagram provides an intuitive means of constructing feedback loops from connectome data.

**Figure 14:**
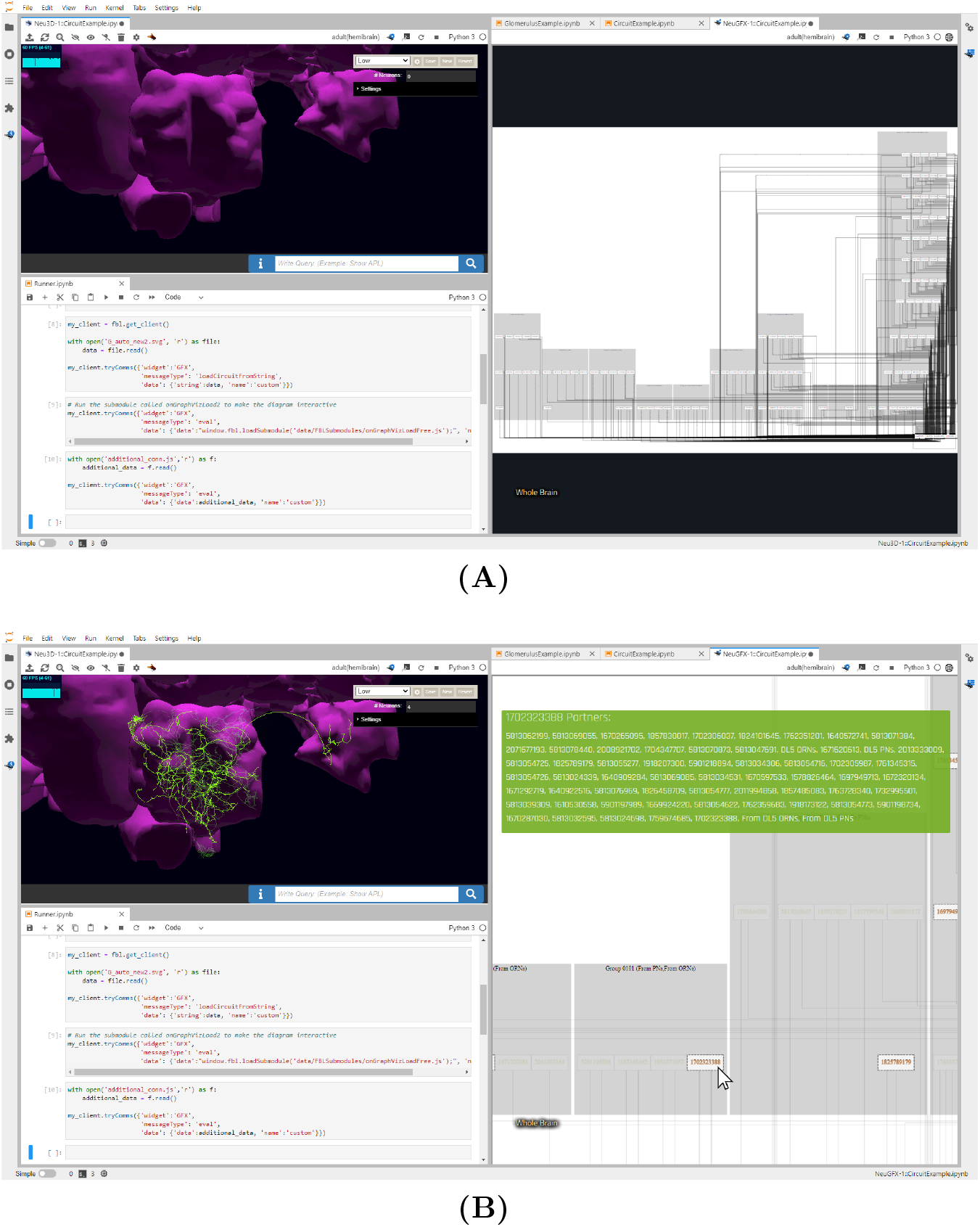
Exploring the feedback loops between the DL5 and DM4 glomeruli using an interactive circuit diagram generated by the FeedbackCircuits Library. **(A)** Users can generate a circuit diagram for any glomerulus consisting of OSNs, PNs, and LNs grouped according to the feedback connectivity pattern described in Figure 8(right) the diagram for the DL5 glomerulus, (top left) the morphology visualization window, (bottom left) the interactive notebook associated with this FlyBrainLab workspace. **(B)** The generated diagrams are interactive. Hovering over the neurons shows their partners, and highlights them in the diagram and in the corresponding 3D morphology; clicking disables/enables them for program execution. Here, the user is currently hovering over the Hemibrain neuron with identifier 1702323388, which shows its partners in the green block on the top right window and highlights it on the morphology in the top left in green.

**Figure 15:**
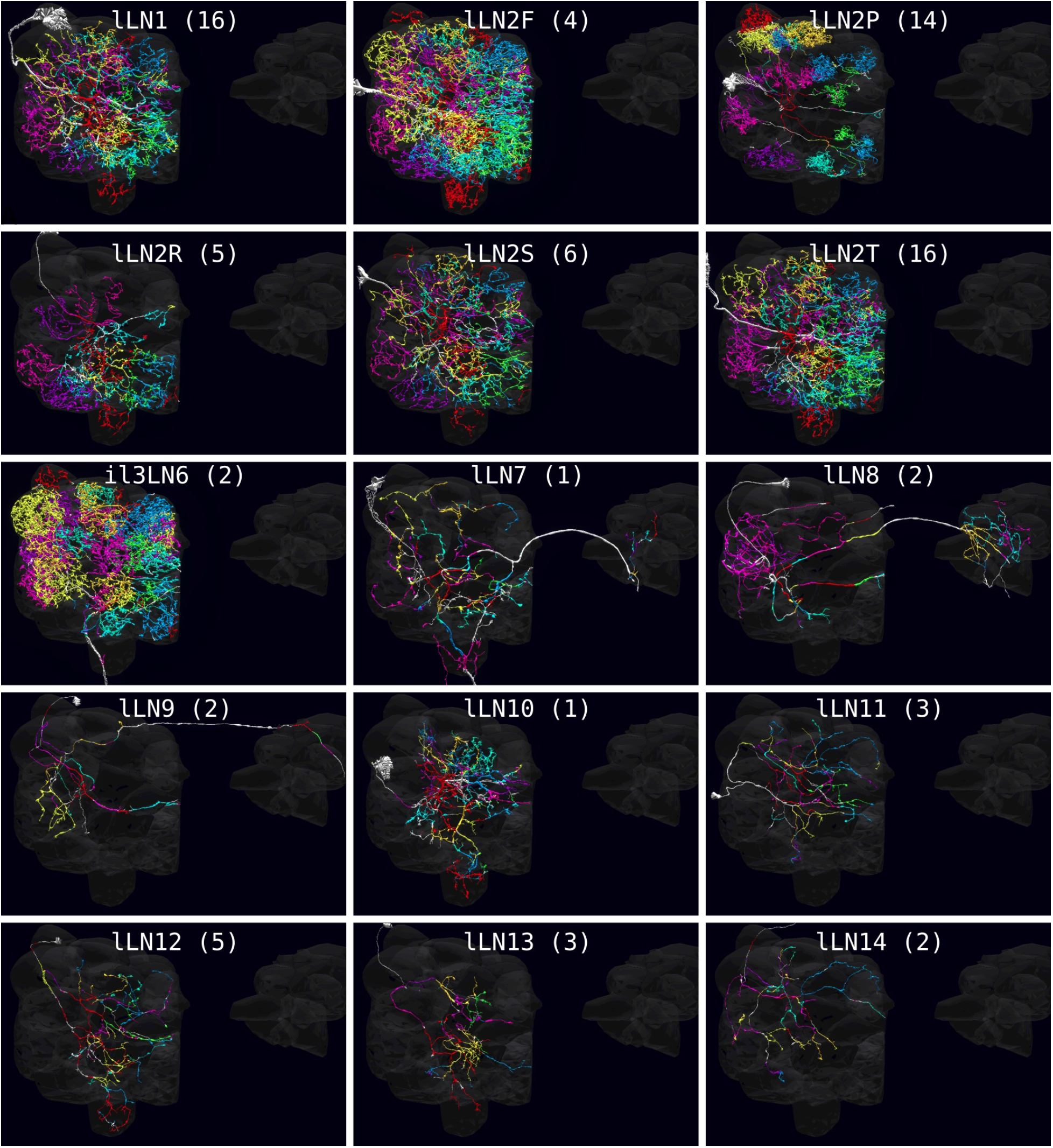
Morphological types of LNs in the Antennal Lobe. An integer in parenthesis indicates the number of neurons of the same cell type.

**Figure 15 (cont.):**
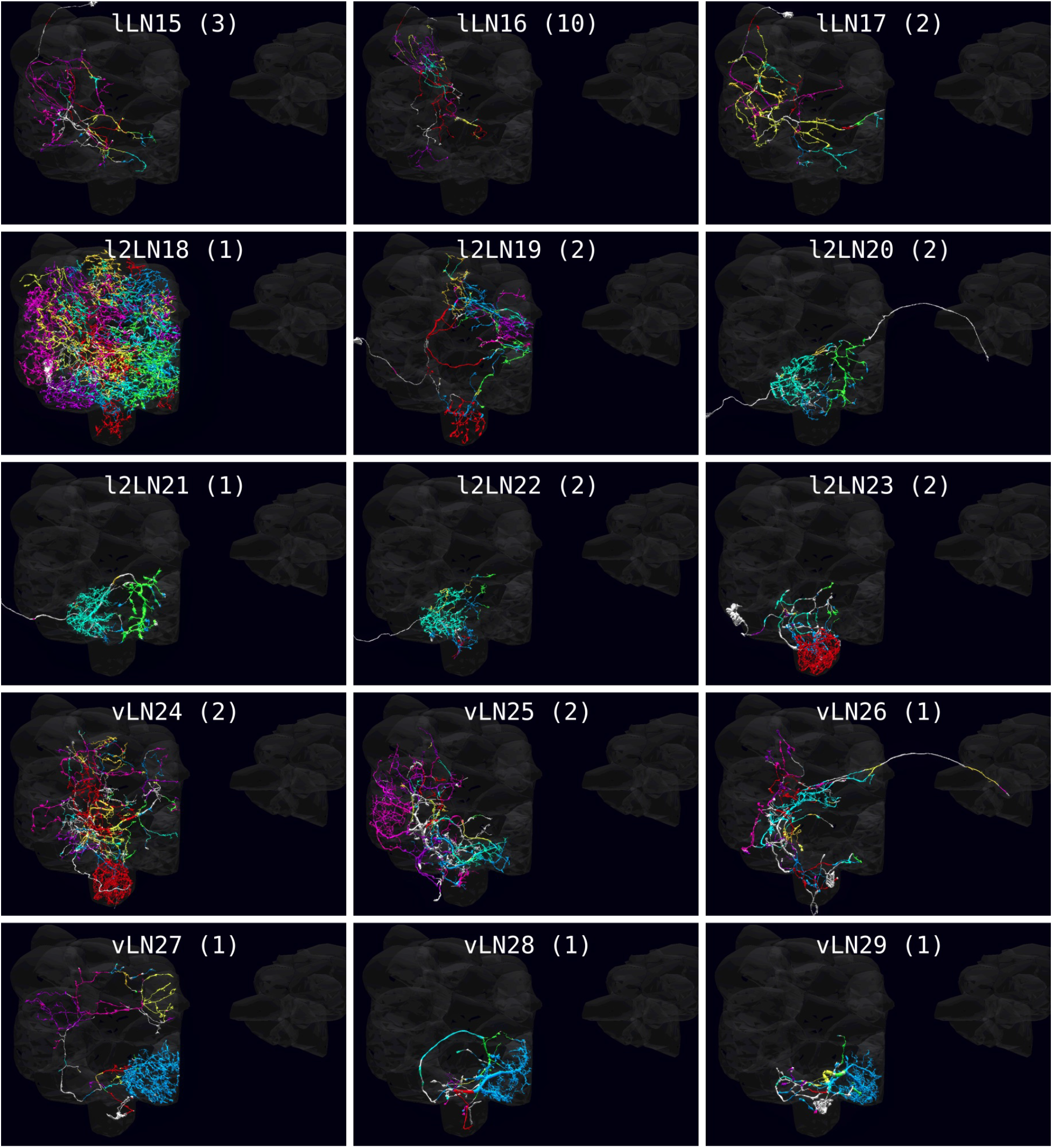
Morphological types of LNs in the Antennal Lobe.

**Figure 15 (cont.):**
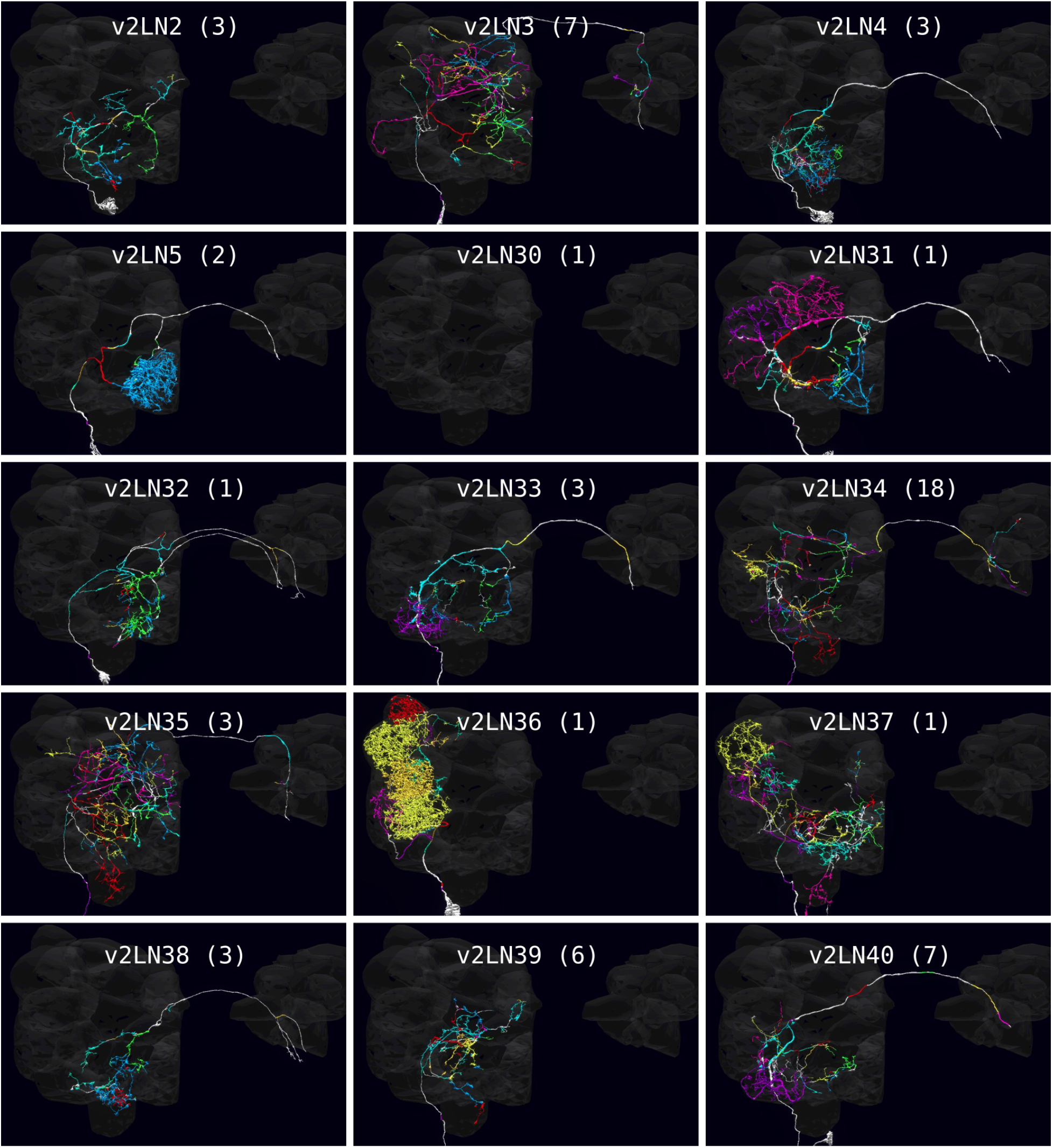
Morphological types of LNs in the Antennal Lobe.

**Figure 15 (cont.):**
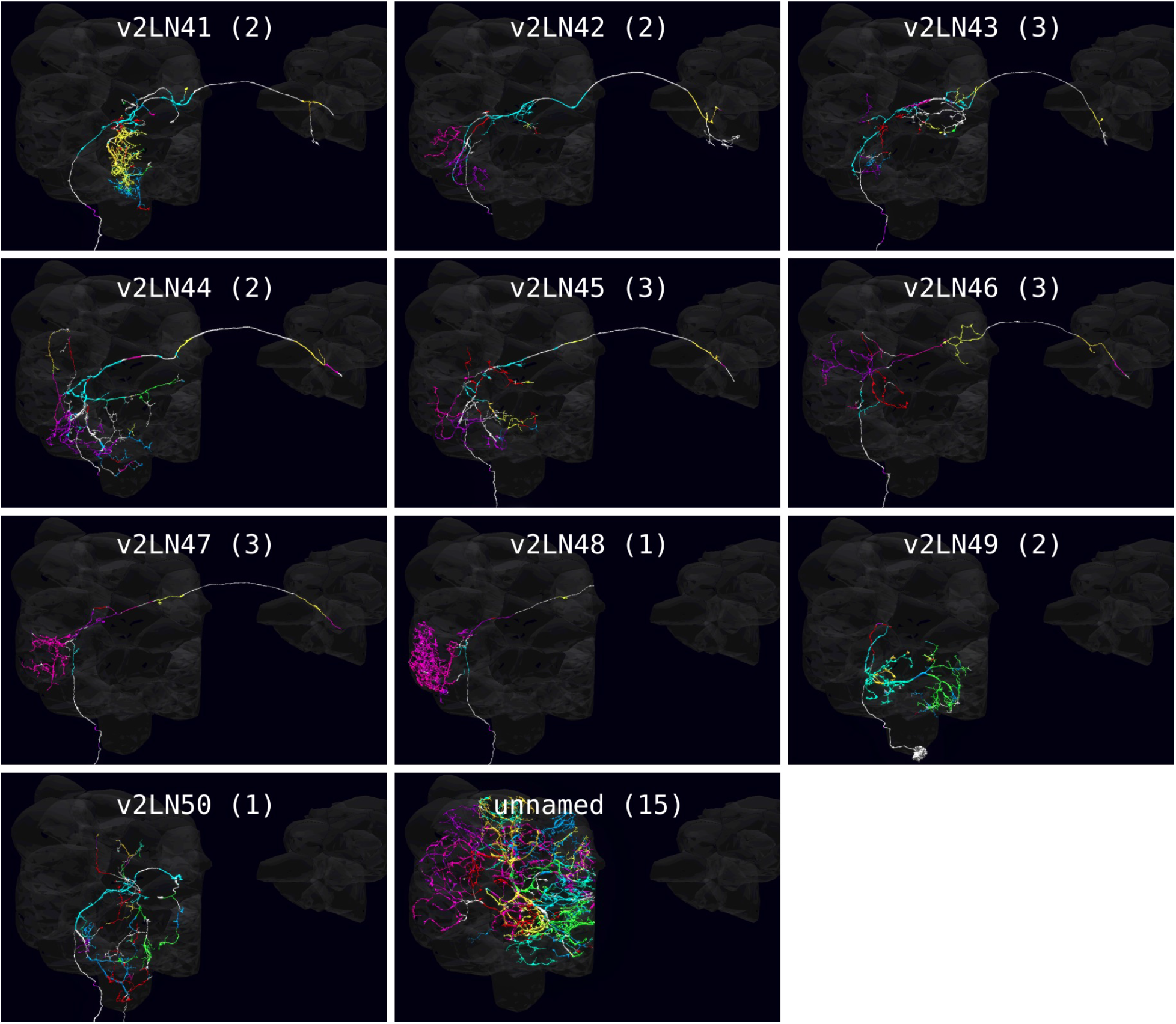
Morphological types of LNs in the Antennal Lobe.

#### Evaluating the Role of Massive Feedback Circuits in a Single Glomerulus

To construct the DM4 glomerulus feedback circuit using the FeedbackCircuits Library, we created a DM4 glomerulus object that includes all the OSNs and PNs and their connections according to the Hemibrain dataset. We then defined three LNs each corresponding to the 3 feedback motifs in Figure 9, and, accordingly connected these LNs to OSNs and PNs. The construction of the DL5 glomerulus feedback circuit follows a similar procedure.

To model the odorant transduction process of the OSNs, we followed the OTP model in [21]. The axon hillock of OSNs, PNs and LNs is given by the Connor-Stevens point neuron model [44]. All synapses are modeled as a variation of the *α* synapse. We also modeled the presynaptic effect of LNs onto the OSN axon terminal. A detailed description of the dynamics of the olfactory transduction process, neurons and synapses can be found in Appendix B.

To evaluate the feedback loop, we sweep through all possible concentration-modulated affinity values, defined as [**a**]_*ron*_ · [**u**]_*ron*_ where [**a**]_*ron*_ is the affinity of receptor *r* with an odorant *o* of OSN *n*, and [**u**]_*ron*_ is a constant concentration waveform of odorant *o* presented to the OSN *n* (see also the x-axes of Figure 11). To interpret the result, one can pick an affinity value, and thus an odorant that interacts with the receptor, and rescale the axis to obtain the response of the PN, projecting into the glomerulus, to different odorant concentration profiles.

#### Evaluating the Role of Massive Feedback Circuits in/between Two Glomeruli

To construct the feedback circuit of two interconnected glomeruli of DM4 and DL5, we start with two independent circuits, each with an LN1 and an LN2, for each of the glomeruli. We then combine these two independent circuits, and add an LN3 that connects to each LN1 and LN2 in both directions. Instead of stimulating LN3 externally, we assume synapses from LN1 and LN2 to LN3 are excitatory.

The olfactory transduction, axon hillock and synaptic models are the same as the ones of the single glomerulus above, and their dynamics are described in detail in Appendix B.

To evaluate the feedback circuit in two interconnected glomeruli, we sweep through constant inputs on a grid of concentration-modulated affinity values associated with the odorant receptor of OSNs that project into the DM4 and DL5 glomeruli, respectively. PN responses to the inputs with values on lines crossing the origin can be used to characterize the responses to the odorants of interest.

## Acknowledgments

The research reported here was supported by AFOSR under grant #FA9550-16-1-0410, DARPA under contract #HR0011-19-9-0035 and NSF under grant #2024607.

## A Antennal Lobe Local Neuron-Types

We present in Figure 15 the full list of LN-types from the NeuroNLP++ query.

## B Neuronal/Synaptic Dynamics and Odorant Transduction Models

We used the Connor-Stevens neuron model [44] whose dynamics can be expressed by the system of differential equations

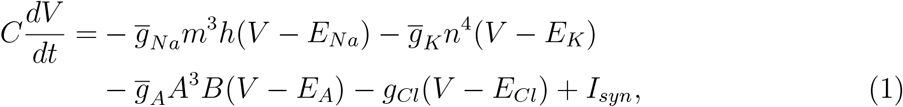

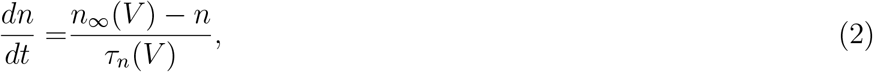

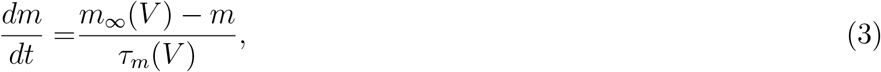

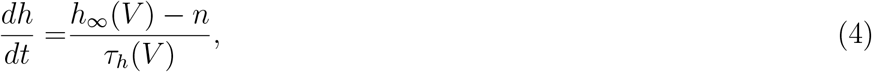

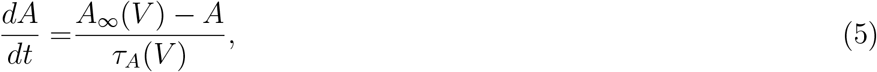

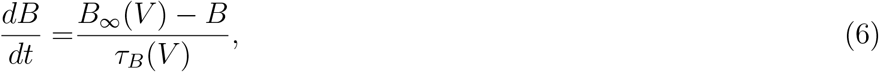

where

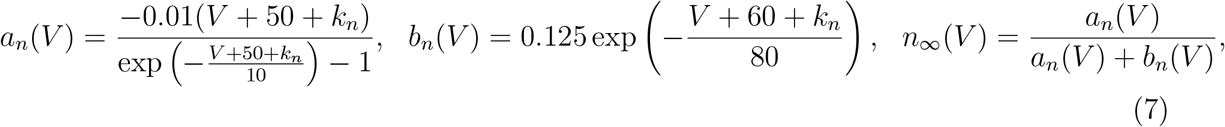

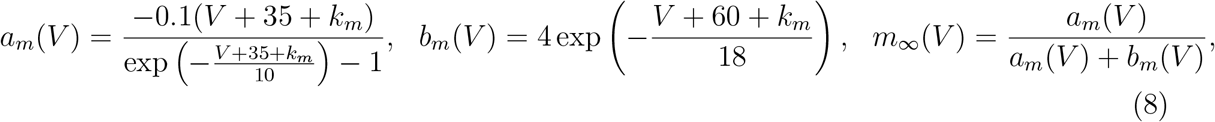

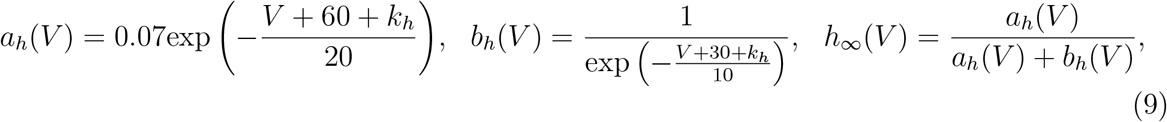

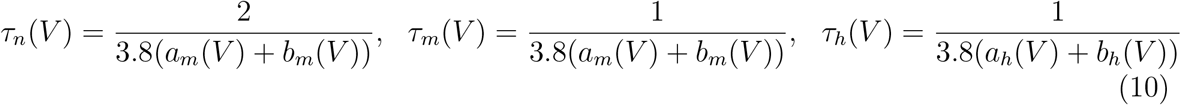

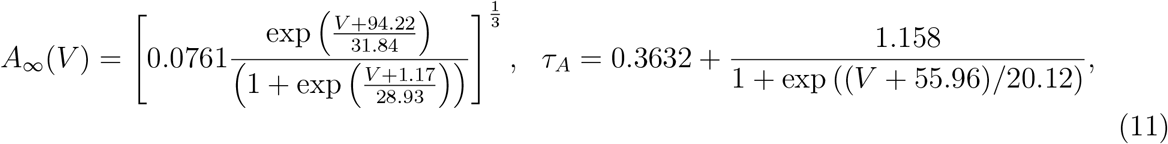

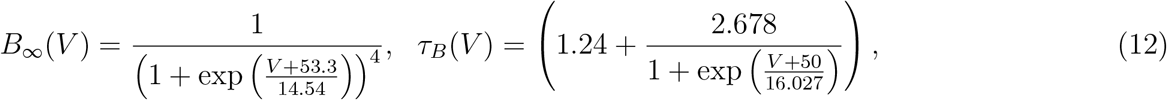

All synapses between OSNs and LNs, LNs and OSNs, OSNs and PNs, LNs and PNs and, PNs and LNs are modeled as *α* synapses described by the following equations:

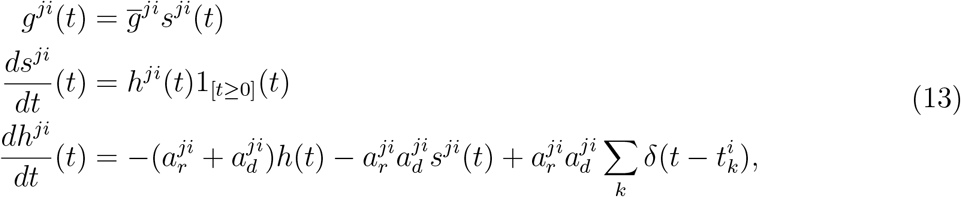

where *i* and *j* are the indices of the presynaptic and postsynaptic neurons, respectively, *s*^*ji*^(*t*) and *h*^*ji*^(*t*) are state variables, and 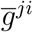 is a scaling factor, 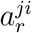 and 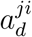 are, respectively, the rise and decay time of the synapse, 1_[*t*≥0]_(*t*) is the Heaviside function and *δ*(*t*) is the Dirac function. 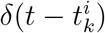 indicates an input spike from the presynaptic neuron at time 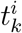.

LN-to-OSN synapses do not provide a current to OSNs. Rather, they act at the presynaptic site on the OSN terminals and affect neurotransmitter release. We model this interaction directly in the postsynaptic current of the OSN-to-PN and OSN-to-LN synapses. The postsynaptic current induced by an OSN-to-PN synapse can be expressed as

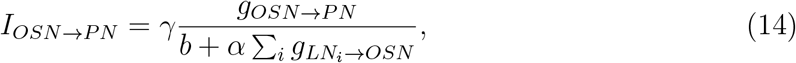

where *b*, *α* and *γ* are constants and *γ* corresponds to a fixed potential of the postsynaptic neuron membrane, and *g*_*OSN*→*PN*_ is the conductance of the OSN-to-PN synapse, and 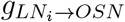 is the conductance of synapses from all the *i*th LN to an OSN. The postsynaptic current induced by an OSN-to-LN synapse shares the same form.

The postsynaptic current induced by an PN-to-LN synapse can be expressed as

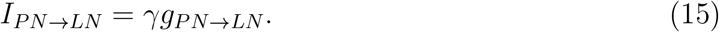

According to [21], the OSN odorant transduction process is given by

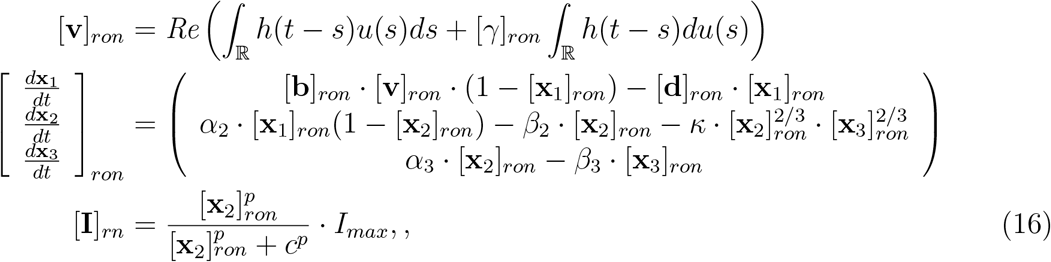

where *o* is the index of a pure odorant, *u*(*t*) is the concentration waveform presented to the antenna, [**I**]_*ron*_ is the transduction current of neuron *n* with receptor *r*. The biological spike generator of the OSN is modeled as a Connor-Stevens neuron with [**I**]_*ron*_ as the current source.

Note that the differential equation on [**x**_1_]_*ron*_ can be rewritten as

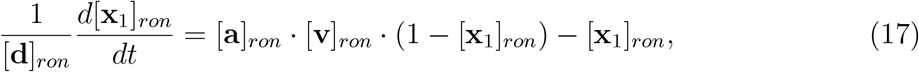

where [**a**]_*ron*_ = [**b**]_*ron*_[**d**]_*ron*_ is the affinity value of the odorant-receptor pair.

